# Passive receptor dissociation driven by porin threading establishes active colicin transport through *Escherichia coli* OmpF

**DOI:** 10.1101/2021.04.15.439832

**Authors:** Marie-Louise R. Francis, Melissa N. Webby, Nicholas G. Housden, Renata Kaminska, Emma Elliston, Boonyaporn Chinthammit, Natalya Lukoyanova, Colin Kleanthous

**Affiliations:** Department of Biochemistry, South Parks Road, University of Oxford, Oxford OX1 3QU, UK; Department of Biological Sciences, ISMB, Birkbeck College, London WC1E 7HX, UK

## Abstract

Bacteria deploy weapons to kill their neighbours during competition for resources and aid survival within microbiomes. Colicins were the first antibacterial system identified yet how these bacteriocins cross the outer membrane of Escherichia coli is unknown. Here, by solving the structures of translocation intermediates and imaging toxin import, we uncover the mechanism by which the Tol-dependent nuclease colicin E9 (ColE9) crosses the outer membrane. We show that threading of ColE9’s disordered domain through two pores of the trimeric porin OmpF causes the colicin to disengage from its primary receptor, BtuB, and reorganise the translocon either side of the membrane. These rearrangements prime the toxin for import through the lumen of a single OmpF subunit, which is driven by the proton motive force-linked TolQ-TolR-TolA-TolB assembly. Our study explains why OmpF is a better translocator than OmpC and reconciles the mechanisms by which Ton- and Tol- dependent bacteriocins cross the bacterial outer membrane.

## Introduction

The asymmetric outer membrane (OM) of Gram-negative bacteria, composed of lipopolysaccharides (LPS) in the outer leaflet and phospholipids in the inner leaflet, shields the organism against environmental insults, the immune systems of plants and animals, bile salts in the human gut and several classes of antibiotics (Nikaido, 2003; Ranf, 2016; Vergalli et al., 2020; Whitfield and Trent, 2014). Bacteria have nevertheless evolved potent antibacterial systems to breach the OM that are integral to the internecine warfare between bacterial populations as they compete for space in which to grow and nutrients on which to feed (Granato et al., 2019). Attack strategies are typically of two types, those that depend on physical or close contact between bacterial cells and those mediated by diffusible molecules (Ruhe et al., 2020). The former includes contact-dependent inhibitors, where a toxin projected from an attacking cell binds a surface receptor of a recipient cell prior to import (Aoki et al., 2010), and type VI secretion, where toxins are delivered by a needle that punctures the OM (Basler et al., 2013). Diffusible toxins are generally referred to as bacteriocins and can be peptides or proteins. Here, we focus on a family of protein bacteriocins from the antimicrobial armoury of the *Enterobacteriaceae* and reveal their convoluted use of porins for OM transport.

Protein bacteriocins (hereafter referred to as bacteriocins) are plasmid or chromosomally-encoded multidomain toxins that kill cells through the action of a C-terminal cytotoxic domain orchestrated by N-terminal regions of the toxin (Kleanthous, 2010). Cytotoxic activity is expressed either in the periplasm, through depolarisation of the cytoplasmic membrane or cleavage of the lipid II peptidoglycan precursor, or in the cytoplasm through cleavage of specific tRNAs, ribosomal RNA or chromosomal DNA. Passage across the OM is catalysed by one of two proton-motive force (PMF)-linked assemblies in the inner membrane. Ton-dependent (group B) bacteriocins contact TonB via TonB-dependent transporters (TBDTs) in the OM whereas Tol-dependent (group A) bacteriocins contact one or more components of the Tol-Pal system (Cascales et al., 2007). Nearly all Tol-dependent bacteriocins identified to-date enter bacteria using a porin in combination with an outer membrane protein receptor, which can be a neighbouring TBDT or the porin itself (Kleanthous, 2010). *E. coli* porins OmpF and OmpC are abundant trimeric β-barrel outer membrane proteins with narrow channels running through them that taper to an eyelet, 7-8 Å at the narrowest point (Nikaido, 2003). The dimensions and electrostatic nature of the eyelet allow the diffusion of nutrients and metabolites (<600 Da) but excludes large antibiotics such as vancomycin and folded proteins.

Recent work has shown that Ton-dependent bacteriocins specific for *Pseudomonas aeruginosa* are transported unfolded through TBDTs (White et al., 2017). Import is energized by the PMF via TonB and its associated ExbB-ExbD stator complex. By contrast, controversy surrounds how Tol-dependent bacteriocins exploit porins to cross the OM and whether this transport is energized. The present work focuses on the group A endonuclease (DNase) colicin, ColE9, a member of the E group of colicins (ColE2-E9), all of which use the same import machinery but target different nucleic acids in the cytoplasm (DNA, rRNA or tRNAs) (Kleanthous, 2010). We demonstrate using cryo-EM and live-cell imaging that ColE9 exploits the trimeric porin OmpF to cross the OM of *E. coli* by a combination of passive and active transport. We show that following binding to its TBDT receptor, BtuB, ColE9 threads its disordered N-terminus through two subunits of OmpF both as a means of capturing a component of the PMF-coupled Tol-Pal system in the periplasm but also to disengage the toxin from its receptor. ColE9 then translocates unfolded through the narrow lumen of a single OmpF subunit by a process that is entirely driven by the PMF. We also demonstrate that large fluorophores normally excluded by the pores of OmpF can be forcibly brought into the cell by attaching them to the colicin.

## Results

### Cryo-EM structure of the ColE9 outer membrane translocon

ColE9 is a plasmid-encoded heterodimer comprising a 60-kDa toxin bound to a small immunity protein, Im9. The two proteins are released together from *E. coli* cells in response to environmental stress (Cooper and James, 1984). Immunity proteins are high-affinity inhibitors that are co-expressed with the colicin and protect the producing cell from its cytotoxic action (Kleanthous et al., 1999), but are jettisoned during import of the bacteriocin (Vankemmelbeke et al., 2013). ColE9 has four functionally annotated domains (**Figure 1a**) (Klein et al., 2016). At the N-terminus, an 83 amino acid intrinsically-unstructured translocation domain (IUTD) houses three protein-protein interaction epitopes (Housden et al., 2010); a 16-residue TolB-binding epitope (TBE) associates with the periplasmic protein TolB and this is flanked by two OmpF-binding sites (OBS1 and OBS2). The IUTD is followed by a folded α/β domain which is thought to be associated with inner membrane translocation and generally referred to as the translocation or T-domain. Following the T-domain is a long coiled-coil receptor-binding (R-) domain the apex of which binds the vitamin B_12_ transporter BtuB on the cell surface (**Figure 1b**). Finally, at the C-terminus of the colicin is a cytotoxic DNase to which the immunity protein Im9 is bound. Following import and ejection of Im9, the internalised DNase elicits cell-death through random cleavages of the bacterial genome. The ColE9 DNase is a member of the H-N-H/ββα-Me class of metal-dependent endonucleases (Pommer et al., 2001), which includes apoptotic nucleases and the core nuclease of CRISPR/Cas9. Other cytotoxic domains can be delivered by the same toxin chassis, including the RNase of ColE3 that inhibits protein synthesis by site-specific cleavage of the ribosomal A-site (Ng et al., 2010). The structures of ColE3 (Soelaiman et al., 2001) and ColE9 (Klein et al., 2016) have been reported previously as have the binary complexes of ColE3 R-domain-BtuB (Kurisu et al., 2003), ColE9 OBS1-OmpF (Housden et al., 2010) and ColE9 TBE-TolB (Loftus et al., 2006).

**Figure 1.**
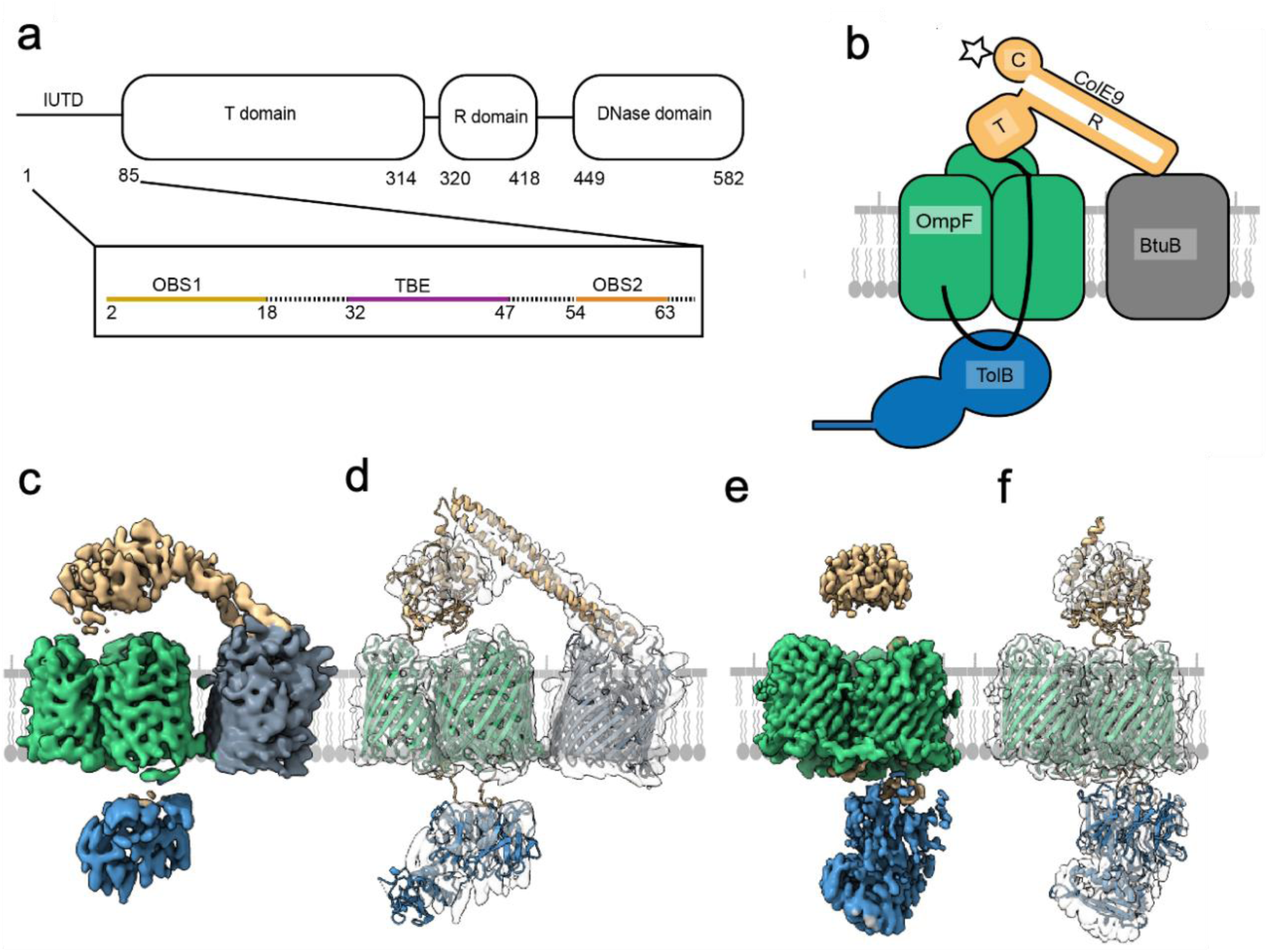
Cryo-EM structures of the ColE9 outer membrane translocon. **a**, Schematic of the ColE9 sequence showing its constituent domains: an intrinsically unstructured translocation domain (IUTD) at the N-terminus is followed by three structured domains involved in translocation (T), receptor (R) binding and cytotoxicity (C). The IUTD houses three linear protein-protein interaction epitopes, two OmpF binding sites (OBS1, OBS2) flank a TolB-binding epitope (TBE). Residue numbers denote position in ColE9 sequence. **b**, Cartoon of the ColE9 OM translocon. ColE9 (*orange*) exploits the vitamin B_12_ transporter BtuB (*grey*) as its extracellular receptor and the porin OmpF (*green*) for threading its N-terminal IUTD (*solid black line*) through to the periplasm where it captures TolB (*blue*). *Star*, represents the site on the ColE9 DNase domain (K469C) where fluorophores were covalently attached throughout this study. **c**, Cryo-EM map of the fully assembled ColE9 translocon, with local resolution range 4.5-16 Å. Component proteins are coloured as in panel b. The structure shows extracellular ColE9 creating a protein bridge between the two OMPs, BtuB and OmpF. The β-barrel of BtuB is tilted by 35° relative to that of OmpF. TolB is located on the periplasmic side of OmpF, with no associations to BtuB. ColE9 and TolB regions of the map have weaker density that those of the β-barrels. **d**, Model of the intact ColE9 translocon generated after docking and rigid-body refinement of individual structures of ColE9 residues 85-580 (PDB ID 5EW5), OmpF (3K19), BtuB (PDB ID 2YSU), and TolB (PBD ID 4JML). **e**, Cryo-EM map of the partial ColE9 translocon with an average resolution of 3.7 Å, which has density consistent with OmpF, TolB, and ColE9 residues 3-314. Map is coloured based on component parts shown in panel b. TolB is much better resolved here than in the full translocon map in panel c, although ColE9 density is weaker. ColE9 and TolB density aligns on the extracellular and periplasmic side of OmpF, respectively. **f**, The refined structure of the partial translocon, generated by docking and refinement of ColE9 residues 85-580 (PDB ID 5EW5), OmpF (PDB ID 3K19), and TolB-ColE9 TBE (PDB ID 4JML). ColE9 residues 3-75 were built *de novo*.

Tol-dependent colicins assemble a translocon at the cell surface that involves its periplasmic target along with the OM receptor and translocator which together establish the import pathway of the toxin. No structures have yet been reported for any colicin translocon and so there is little understanding of how these assemblies underpin OM transport. We have reported previously the isolation and purification of the entire OM translocon for ColE9 in which the toxin, bound to TolB, BtuB, and OmpF, is solubilised in 1% β-octylglucoside (β-OG) (Housden et al., 2013). This complex is in addition stabilised by a disulphide bond at the periphery of the ColE9 TBE-TolB interface which does not impact the protein-protein interface (Housden et al., 2013). We followed the same procedure, which involves a combination of *in vivo* and *in vitro* approaches, but, for ease of assembly (see Materials and Methods), using ColE9 truncated at the end of the R-domain and hence missing both the DNase and Im9. The assembled translocon was transferred into amphipols and vitrified on graphene oxide coated grids for visualisation of individual particles by cryo-EM (see Materials & Methods for details).

Single particle analysis revealed the presence of two populations of ColE9 translocon complexes, one where the translocon was intact and a second class where BtuB was absent, which we refer to as the partial translocon (**Supplementary Figures 1 and 2**). The final post-processed map of the full translocon (**Figures 1c and d**) is at an overall resolution of 4.7 Å, according to the gold-standard FSC method, with local resolutions ranging from 4.7 to 16 Å. The absence of BtuB in the partial translocon resulted in an improvement in map resolution (**Figures 1e and f**), which varies from 3.7 to 6 Å (FSC). Masked subtraction of BtuB from the full translocon map and refinement of these subtracted particles did not improve map quality, suggesting that it is the loss of BtuB from the complex which resulted in the increase in map resolution. Density for ColE9 was better resolved in the full compared to the partial translocon where only the T-domain was resolved. This is most likely due to the ColE9 R-domain providing some conformational rigidity in the fully assembled translocon through its interaction with BtuB. This contention is supported by the loss of density for the R-domain in the partial translocon map. By contrast, density for TolB was better resolved in the partial translocon map. The partial translocon map presented here is similar to a negative stain map of a proteolysed version of the translocon (comprising of OmpF, TolB and ColE9 residues 1-122), reported previously at a resolution ∼20 Å ((Housden et al., 2013). However, the lower resolution of this earlier structure meant little information was forthcoming about ColE9 interactions with OmpF.

Isolated crystallographic structures for the majority of ColE9 translocon components provided a starting point for model building into the full and partial translocon maps. Although the resolution of the full translocon map did not allow for local refinement of docked structures, rigid-body refinement was carried out to generate a molecular model. For the partial translocon model, OmpF was docked and refined locally along with the ColE9 IUTD (residues 3-75), which were built *de novo* into the map. The remaining ColE9 T-domain residues (85-316) and TolB were rigid-body refined into the partial translocon map due to the lower resolution associated with these parts of the map.

In the fully assembled translocon model, the 22-stranded β-barrel of BtuB sits at a 35° angle relative to the trimeric β-barrel structure of OmpF (**Figures 1c and d**). Unmodelled density, most likely that of a single molecule of LPS, is wedged between the two proteins which likely contributes to displacement of BtuB relative to OmpF. Native state mass spectrometry has shown previously that a single LPS molecule remains associated with the purified translocon complex (Housden et al., 2013). Since there are no examples in the PDB of heterologous bacterial outer membrane proteins residing next to each other it is not clear if the 35° tilt reflects the natural position of BtuB in the OM or the way in which the ColE9 translocon was assembled for cryo-EM analysis. We note however that if the β-barrels of OmpF and BtuB were flush in the membrane the 45° trajectory of the ColE9 R- domain-BtuB complex, first shown for the ColE3 R-domain-BtuB complex (0.95 Å rmsd) (Kurisu et al., 2003), would project the T-domain beyond OmpF. The coiled-coil nature of the R-domain, with BtuB at its apex, means the cytotoxic DNase domain although absent from the current structure would sit ∼60 Å above OmpF. Finally, TolB is held on the periplasmic side of OmpF through its association with the ColE9 TBE, but is not well-resolved in the full translocon model (**Figures 1c and d**).

The partial translocon is comprised of OmpF, the IUTD and T-domain of ColE9 and TolB (**Figures 1e and f**). Previous work on the translocation mechanism of group A colicins has demonstrated that these colicins contact their periplasmic binding partners (typically TolB and/or TolA) by a direct epitope delivery mechanism following binding to the cell surface receptor (Housden et al., 2010; Jansen et al., 2020). The colicin threads its IUTD through two of OmpF’s three pores thereby presenting a tethered protein-protein interaction epitope to the periplasm. In the case of ColE9, the TBE promotes contact between TolB and TolA in the inner membrane which is coupled to the PMF through its partners, TolQ and TolR (Bonsor et al., 2009). The partial translocon structure of ColE9 shows how the IUTD (residues 3-75) navigates through the lumen of one OmpF subunit in order to thread back up into a neighbouring subunit. The end result is that each OBS is lodged within an individual subunit of OmpF: OBS1 (residues 2-18) is bound to subunit 1 while OBS2 (residues 54-63) is bound to subunit 2, leaving subunit 3 free of colicin. The relative positions of the two OBSs in the partial translocon model implies that subunit 2 is the entry port for the ColE9 IUTD following its initial binding to BtuB in the OM.

The resolution of the full translocon does not allow for detailed comparison of OBS binding with that of the partial translocon (details below), however an overlay of the two models reveal that the ColE9 T-domain undergoes a large-scale movement, rotating by 15° and moving vertically by 12 Å along the rotation axis (**Figures 2a-c**). In the full translocon, the orientation of the ColE9 T-domain is fixed by the R-domain’s contact with BtuB. Rotation of the ColE9 T-domain in the partial translocon to achieve the orientation adopted in the full translocon structure results in a poorer fit to map density. Importantly, rotation of the ColE9 T-domain from the full to the partial translocon repositions ColE9 from a central position above the OmpF trimer to become localised above subunit 2 within which OBS2 is bound (**Figures 3a and c**).

**Figure 2.**
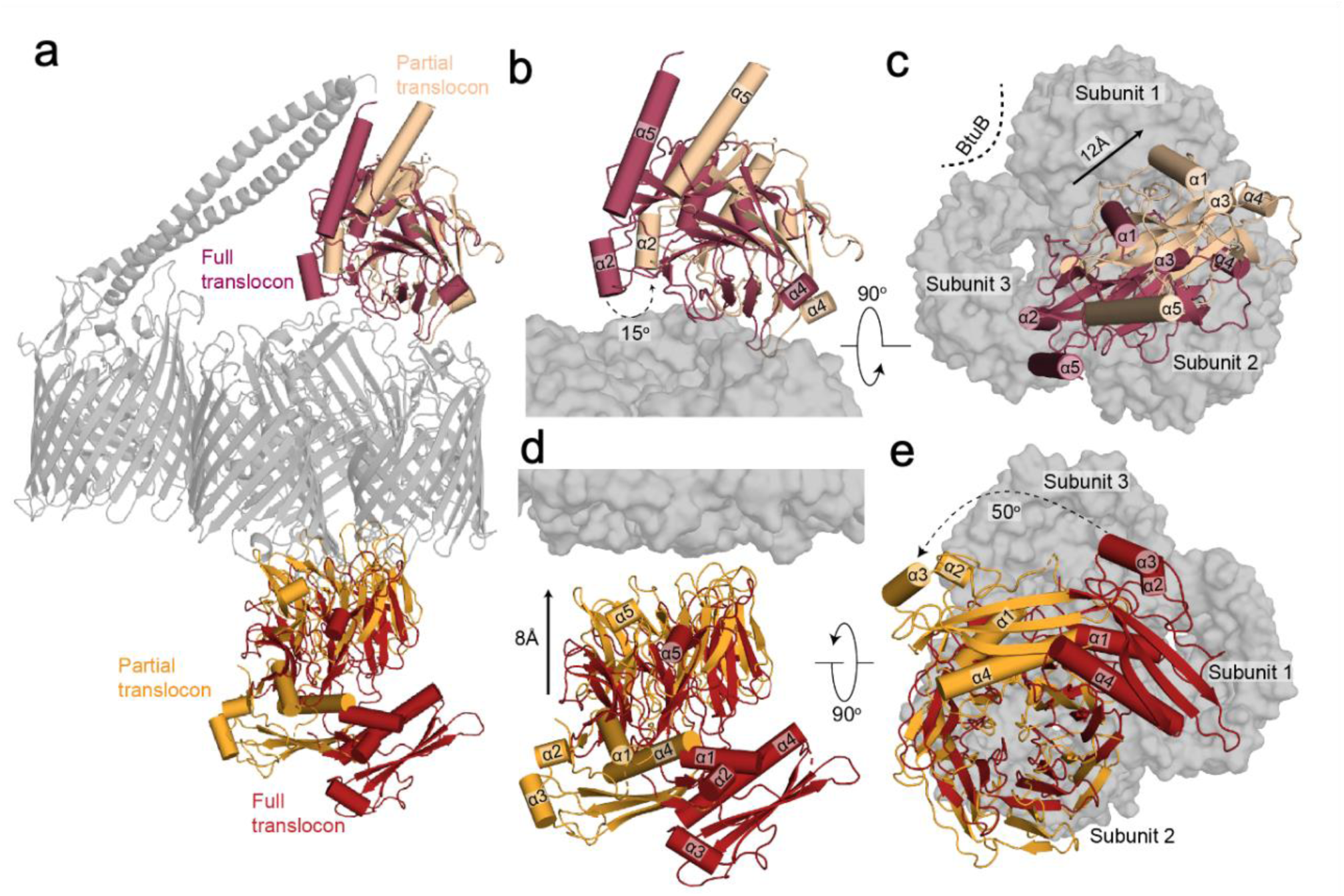
Large-scale structural rearrangements accompany the loss of BtuB from the ColE9 translocon. **a,** Superposition of the complete and partial ColE9 translocon structures (*grey*) aligned on OmpF. TolB in *red* and *orange* denote the full and partial translocons, respectively, ColE9 T-domain is presented in *crimson* and *pale orange* for the full and partial translocons, respectively. **b**, Sideview comparison showing the relative positions of the ColE9 T-domain (residues 85-316) in the two structures and highlighting the 15° rotation that occurs transitioning from the full (*crimson*) to the partial (*pale orange*) translocon. **c**, Extracellular view of the ColE9 T-domain position, with the OmpF trimer shown in the background. The loss of BtuB from the translocon complex elicits a 12 Å movement along the axis of rotation (*black arrow*) that results in repositioning of the T-domain from a central location (*crimson*) to above subunit two of OmpF (*pale orange*). **d**, TolB undergoes both rotation and translation when transitioning from the full (*red*) to the partial translocon (*orange*). The C-terminal β-propeller domain of TolB, which binds the ColE9 TBE, moves toward OmpF in the OM by ∼8 Å along the rotation axis. **e**, View along the rotation axis from the periplasmic side of OmpF (*grey surface*) highlighting the 50° rotation that TolB undergoes upon loss of BtuB from the translocon complex.

**Figure 3.**
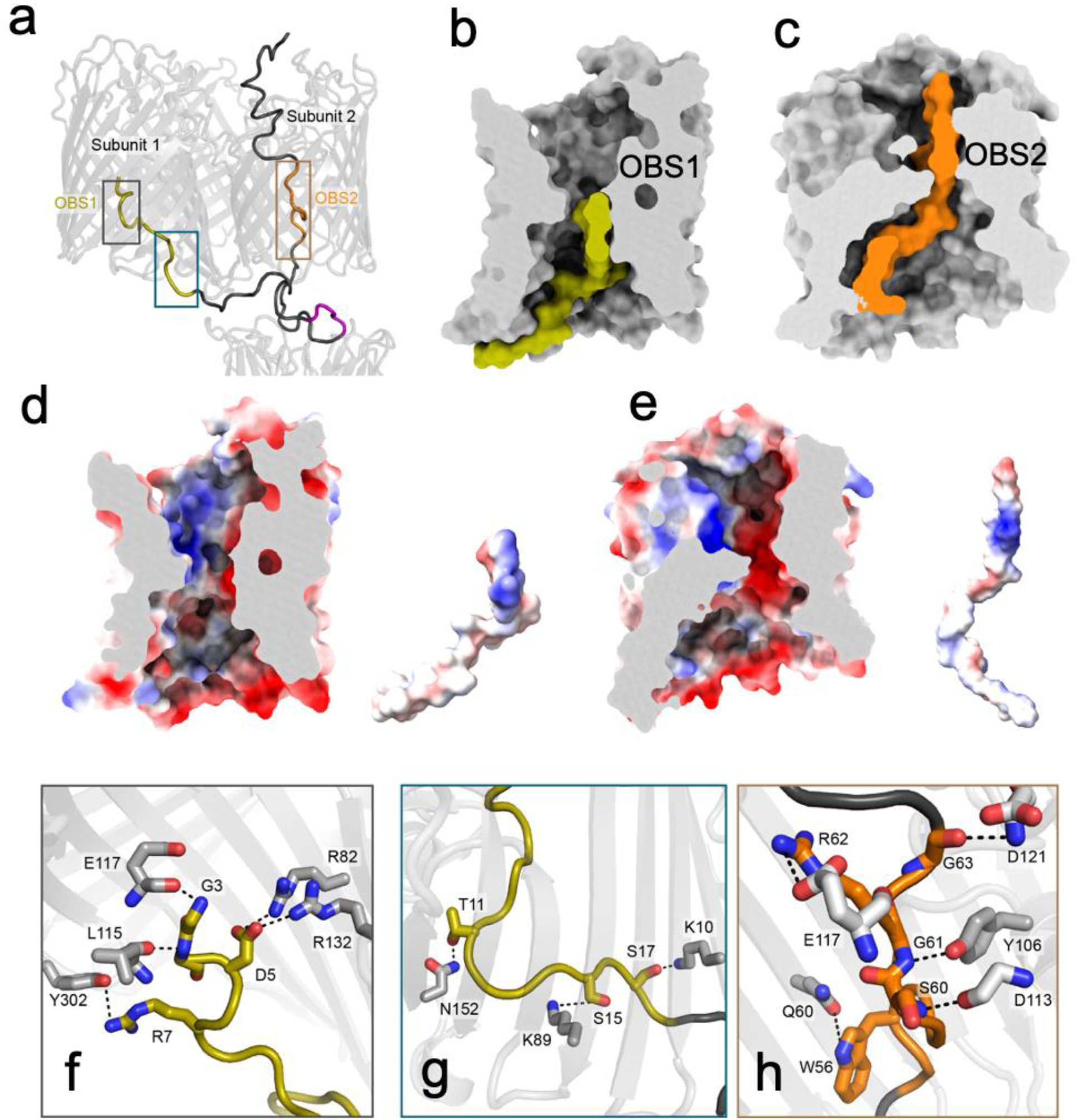
Distinct modes of ColE9 OBS1 and OBS2 recognition within the pores of OmpF enable threading and the tethered presentation of the TBE to the periplasm. **a**, ColE9 residues 3-75 of the IUTD in the partial translocon structure pass through OmpF subunit 2 and then back up into subunit 1. As a result, OBS1 (*gold*) docks within subunit 1 and OBS2 (*orange*) within subunit 2. The ColE9 TBE motif (*pink*) interacts with TolB is positioned below subunit 2, above which the ColE9 T-domain is located. **b**, Surface representation of OmpF subunit 1 (*grey*) and ColE9 residues 3-24 (*gold*) containing OBS1 sequence. ColE9 OBS1 binds the inner vestibule of OmpF subunit 1 such that it runs along the edge of the vestibule, stopping at the eyelet. **c**, ColE9 residues 45-70 (*orange*) encompassing OBS2, traverse the pore of OmpF subunit 2 (*grey*). After passing through the eyelet, OBS2 tracks more centrally into the inner vestibule unlike in panel c where residues 3-24 trail the side of the pore. **d**, OmpF subunit 1 displayed as an electrostatic surface (same cut-through as in b), revealing a patch of negative charge located on the extracellular side of the eyelet that interacts with OBS1, which is contained within the sequence (residues 3-24) also shown as an electrostatic surface adjacent to the subunit. **e**, OmpF subunit 2 displayed as an electrostatic surface (same cut-through as in c), with residues 45-70 also shown as an electrostatic surface adjacent to the subunit. Projecting out of the page in the OBS2 sequence is a positively charged region flanked by negative charges. **f**, A zoom-in of grey box in panel a highlighting hydrogen bonding network between residues 3-7 of OBS1 (*gold stick*s) and nearby residues within OmpF subunit 1 (*grey sticks*). **g**, OBS1 residues 11-17 (*gold sticks*) also interact with the base of OmpF subunit 1 (*grey sticks*) in the periplasm. Region is a zoom-in of blue box in panel a. **h**, OmpF subunit 2 residues (*grey sticks*) form a hydrogen bond network with the backbone of residues of ColE9 OBS2. Region is a zoom-in of brown box in panel a. All electrostatic surfaces shown in panels d and e were calculated using the APBS plugin within Pymol.

TolB also undergoes large-scale movements when transitioning from the full to partial translocon structures, which are likely linked to the re-arrangement of the ColE9 T-domain on the extracellular side of the porin (**Figures 2d and e**). The position of TolB is rotated by ∼50° between the two structures and in the partial translocon is located closer to OmpF through a ∼8 Å upward movement along the axis of rotation. The positioning of TolB closer to OmpF in the partial translocon structure is stabilised by additional interactions between the periplasmic region of OmpF subunit 1 and the IUTD sequence immediately preceding the TBE (**Figure 3g**). Irrespective of the observed movements between the two Cryo-EM models, TolB remains located beneath subunit 2 within which OBS2 is bound.

In summary, we have solved the first cryo-EM structure of a colicin OM translocon. The structure of ColE9 is associated with its full complement of OMP translocation components, BtuB and OmpF, as well as its periplasmic target TolB. We have also solved a partial translocon structure in which the primary receptor BtuB is missing, which leads to substantial rearrangements of ColE9 and TolB above and below the membrane, respectively. As we demonstrate below, these structural changes within the translocon complex are integral to the mechanism by which ColE9 translocates across the OM.

### ColE9 IUTD interactions with the subunits of OmpF

The higher resolution structure of the partial ColE9 translocon provides a molecular explanation as to how bacteriocins exploit porins to establish their translocon complexes at the OM. The lumen of an OmpF monomer is not a straight channel but akin to an hour-glass (**Figures 3a-c**). The outer (extracellular side) and inner (periplasmic side) vestibules of OmpF have diameters of 25 Å and 30 Å, respectively, constricting at the eyelet of the channel to ∼7 Å. After the IUTD region passes through OmpF subunit 2, OBS1 threads back up into subunit 1 such that it snakes along the bottom of the lumen at its widest point and finishes up at the eyelet (**Figures 3a and b**). By contrast, OBS2 spans the eyelet of OmpF subunit 2 via a kink created at Gly61, which is enforced by the hydrogen bonding of neighbouring residues, Ser60 and Arg62 (**Figure 3c**). After OBS2 traverses the eyelet, the remaining ColE9 sequence does not track the OmpF interior as observed for OBS1, instead remaining more centrally located in the inner vestibule.

The bound conformations of the two OBSs are stabilised by electrostatic and hydrogen bonding interactions (**Figures 3d-h**). The vestibules and eyelet of OmpF are highly charged environments; the porin exhibits slight cation selectivity in solute diffusion studies (Basle et al., 2006). Charged patches within the eyelet of OmpF are matched by opposing charges of the OBS sequences (**Figures 3d and e**). In addition to these interactions, networks of hydrogen-bonds lock OBS1 and OBS2 of ColE9 into defined conformations within their respective OmpF subunits (**Figures 3f and h**). Key OBS1 residues that interact with subunit 1 include Asp5, Arg7, Thr11, Ser15, and Ser17 (**Figures 3f and g**). Previous studies have shown that mutation of Asp5, Arg7 or Thr11 to alanine substantially weakens OmpF binding by OBS1 while an Asp5Ala/Arg7Ala double mutant abolishes OmpF binding, consistent with their essential role in stabilising the OBS1-OmpF subunit 1 complex (Housden et al., 2005; Housden et al., 2010). We have reported previously the 3 Å crystal structure of an OBS1 peptide bound to OmpF (Housden et al., 2010). At the time, there was uncertainty as to the orientation of the peptide, which was refined with the N-terminus facing the periplasm. We re-processed these data and rebuilt the model based on the orientation observed in the cryo-EM data for the partial translocon, validating this re-processed model (**Supplementary figures 2k-m**). The new model shows conclusively that OBS1 has its N-terminus pointing towards the extracellular environment (**Figure 3a**) and in both models the conformation adopted by OBS1 is identical. This orientation agrees with recent electrophysiological data and live-cell imaging experiments all of which show OBS1 binding subunit 1 of OmpF from the periplasm (Housden et al., 2018; Lee et al., 2020). In contrast to OBS1, the hydrogen bond network that stabilises OBS2 within subunit 2 of OmpF primarily involves OBS2 backbone atoms and OmpF side-chains (**Figure 3h**), likely explained by the high glycine content within the OBS2 sequence (6/11 residues are glycine, compared to 6/16 for OBS1). The high glycine content also explains how OBS2 can bind in an extended conformation to OmpF, traversing the narrow eyelet. Only two OBS2 residues, Trp56 and Arg62, form side-chain hydrogen bonds with OmpF.

In conclusion, the bound conformations of the OBS epitopes within the electrostatic pores of OmpF are stabilised by salt bridges, backbone and sidechain hydrogen bonds and aided by their high glycine content. The specific binding modes of each OBS is explained by their specific set of interactions within identical OmpF pores and, in the case of OBS1, through directional insertion into the binding site from the periplasm.

### Threading ColE9’s IUTD through the pores of OmpF passively displaces the toxin from its primary receptor, BtuB

The structural differences evident between the full and partial ColE9 translocons centre around the loss of BtuB from the complex. Concomitant with this loss is the restructuring of the ColE9 T-domain and TolB so that, linked by the OBS2 epitope, the two proteins become positioned immediately above and below, respectively, subunit 2 of OmpF. Given the precise nature of these rearrangements we hypothesised that the partial translocon may be indicative of an intermediate state subsequent to ColE9 docking onto the cell surface. To test this hypothesis, we exploited the observation that the isolated R-domain of ColE9 inhibits the growth of an *E. coli* strain dependent on vitamin B_12_ (**Figure 4a**) (Penfold et al., 2000). *E. coli* 113/3 is a *metE* mutant that requires vitamin B_12_ to synthesise methionine. Since the 76-residue ColE9 R-domain competitively blocks binding of vitamin B_12_ to BtuB and binds with higher affinity its addition to *E. coli* 113/3 grown in minimal media supplemented with vitamin B_12_ suppresses cell growth. We repeated this experiment using an identical concentration of intact ColE9 harbouring a single mutation, W39A, within its TBE. Substitution of ColE9 Trp39 for alanine inactivates the toxin by abolishing binding to TolB in the periplasm, which in turn disengages the colicin from the PMF-coupled TolQ-TolR-TolA complex in the IM (Hands et al., 2005). Hence, while all other OM interactions of the colicin are those of the wild-type toxin, ColE9 W39A cannot deliver its DNase into the cell. Addition of ColE9 W39A to *E. coli* 113/3 largely restored cell growth under conditions where the isolated R-domain inhibited growth (**Figure 4a-c**). These data imply, along with the cryo-EM data, that subsequent to binding BtuB in the OM, threading of the ColE9 IUTD through two subunits of OmpF to form the translocon results in the toxin disengaging from BtuB. This translocon-induced dissociation is a passive process since ColE9 W39A is unable to engage with the energised Tol-Pal system. We next investigated if and when the PMF is engaged in OM transport of the colicin.

**Figure 4.**
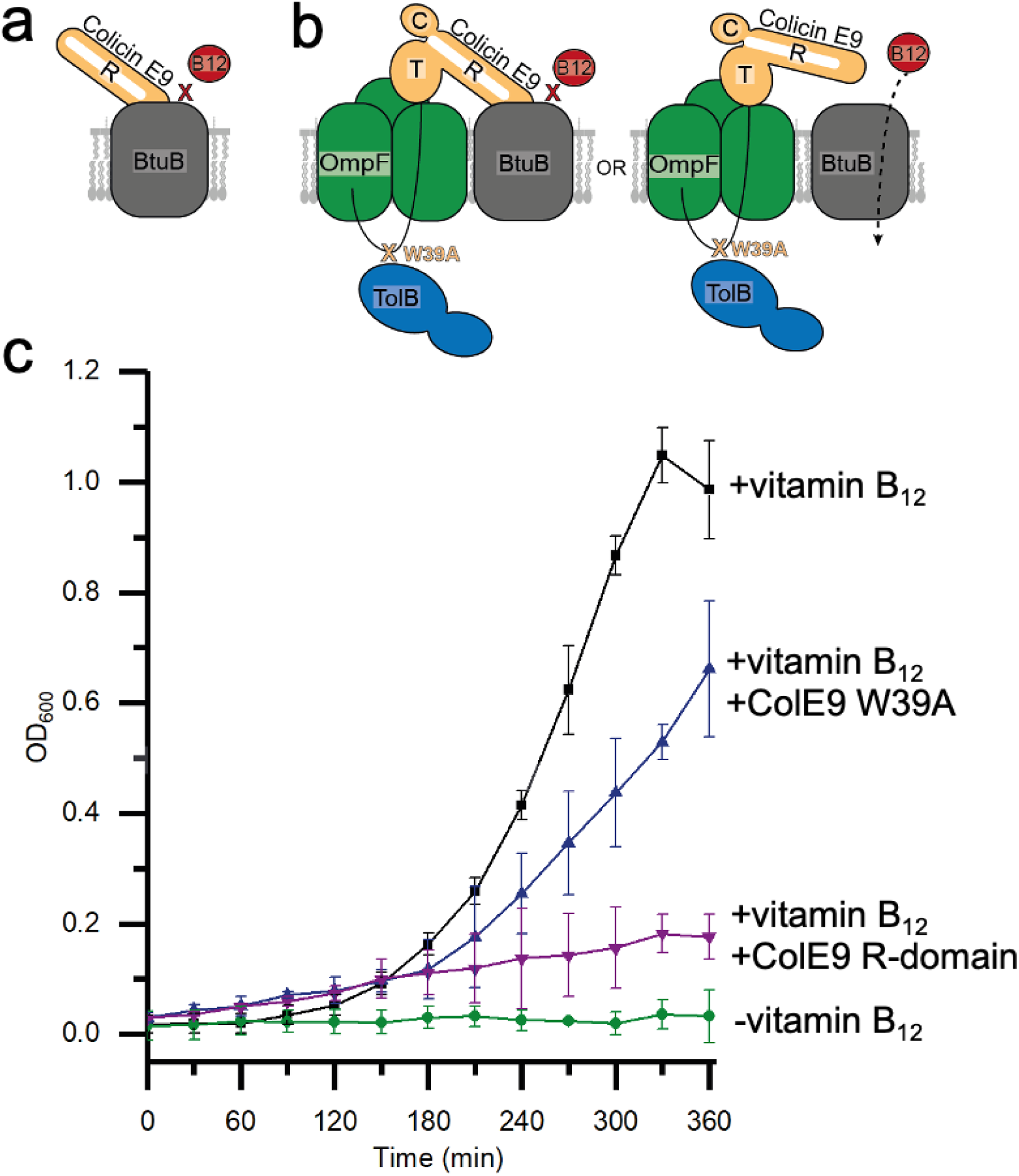
ColE9 threading through the subunits of OmpF disengages the toxin from its outer membrane receptor, BtuB. **a**, Cartoon depicting the basis for the control where the isolated R-domain of ColE9 (residues 348-418, *orange*), which binds BtuB (*grey*) with higher affinity than vitamin B_12_ (*red*), impairs growth of *E. coli* 113/3, a B_12_-dependent strain (Penfold et al., 2000). **b**, Cartoon schematic of the two possible translocon outcomes for ColE9 W39A and their implications on vitamin B_12_ import in *E. coli* 113/3. The W39A mutation within the TBE of ColE9 (*orange*) impairs TolB (*blue*) binding thereby uncoupling the OM components of the translocon from the energized TolQ-TolR-TolA complex in the inner membrane. If threading of the ColE9 IUTD through OmpF (*green*) has no impact on ColE9 binding to BtuB (*grey*) then vitamin B_12_ (*red*) cannot enter cells to support growth. In contrast, if OmpF threading disengages ColE9 from BtuB then vitamin B_12_ can enter cells to support growth. **c**, Growth curves for *E. coli* 113/3 in defined media and in the presence or absence of B_12_ (*black* and *green*, respectively). When challenged with 40 nM isolated R-domain (*purple*), cell growth was largely impaired whereas growth in the presence of 40 nM ColE9 W39A (*blue*) approached that of the no colicin control, consistent with OmpF threading causing the disengagement of ColE9 from BtuB.

### The electrical potential of the PMF drives ColE9 import across the OM

Assembly of ColE9’s OM translocon occurs without the input of cellular energy, including dissociation from the BtuB receptor. At this point in the import process ColE9 is composed of multiple folded domains so this raised the question of whether transport across the OM involved the PMF in order that domain unfolding could occur. Earlier studies suggested some form of unfolding to be involved; inferred from the activating effects of urea on colicin toxicity and the inhibitory effects of intramolecular disulphide bonds (Benedetti et al., 1992; Housden et al., 2005). However, the involvement of the PMF in Tol-dependent nuclease colicin entry is controversial. A recent model, for example, suggests that instead of energised transport, passive, electrostatically-driven ratcheting of the colicin nuclease occurs through the one remaining unoccupied subunit of OmpF (subunit 3 in the present work) once the translocon is formed (Cramer et al., 2018).

A major issue the bacteriocin field has faced thus far in establishing if transport across the OM is energised is the lack of appropriate measures of transport that are not dependent on cell death as a readout. To address this problem, we developed a live-cell imaging platform using fluorescently-labelled ColE9. A single cysteine mutation was engineered at Lys469 in the ColE9 DNase domain (**Figure 1b**), which would be the last domain of the colicin to enter if the toxin was pulled into the cell by the Tol-Pal system. Lysine 469 is adjacent to the DNase active site (**Supplementary figure 3a**). The mutant was labelled initially with AlexaFluor (AF) 647 and excess dye removed by gel-filtration chromatography. Labelling efficiency was typically 90-100% for this and all other fluorophores used in this study (see Materials and Methods for details). The impact of labelling ColE9 K469C on biological activity was assessed by a plate-killing assay, where serial dilutions of the colicin were pipetted onto a lawn of susceptible *E. coli* JM83 cells and grown overnight (**Supplementary figure 4**). Wild-type ColE9 typically shows killing activity against *E. coli* JM83 (a K-12 strain) down to ∼50 pM. Labelling ColE9 K469C with AF647 reduced colicin activity by ∼two-orders of magnitude relative to wild-type ColE9, which nevertheless represents significant toxicity since clearance zones are visible at low nanomolar concentrations. The decrease in activity could be the result of the fluorophore affecting cytotoxicity and/or import. Alternatively, if labelling completely abolished colicin activity the residual toxicity might simply reflect the presence of unlabelled molecules.

In order to distinguish between these possibilities, we investigated the effect of labelling the same position with fluorophores of differing size and charge (AF488, AF568, AF647). We rationalized that if decreased colicin activity reflected unlabelled molecules then altering the fluorophore would have a negligible effect given that the maleimide chemistry used to attach the fluorophores was the same in each case. None of the fluorophores attached at this position affected the ability of the isolated nuclease domain of ColE9 to degrade dsDNA (**Supplementary figure 3b-e**). We next repeated the *E. coli* JM83 cell-killing assay for the ColE9 K469C mutant labelled with each of the fluorophores. These experiments showed that the fluorophores affected toxicity to varying degrees; the order of their impact being AF568>AF647>AF488 (**Supplementary figure 4**). Indeed, ColE9 K469C^AF568^ displayed very little colicin activity. Finally, we conducted further *in vivo* tests by assessing the impact of fluorophore labelling ColE9 K469C on the activation of the DNA damage-induced SOS response in *E. coli* cells (**Supplementary figure 5a-d**). Transported ColE9 readily activates SOS through cleavage of chromosomal DNA, which can be monitored by *lux* bioluminescence in an *E. coli* reporter strain (DPD1718) (see Materials and Methods). As reported previously (Vankemmelbeke et al., 2005), ColE9 above 20 nM completely arrested growth of DPD1718 in liquid culture at 37 °C after 90 min and this corresponded to a peak in *lux* bioluminescence, which then subsided to zero as all cells were killed-off (**Supplementary figure 5b)**. Lower concentrations of ColE9 resulted in the same initial increase in *lux* bioluminescence but did not subside because cell growth begins to outcompete colicin toxicity. While none of the fluorophore labelled ColE9 constructs (at 20 nM) significantly impacted growth of *E. coli* DPD1718, all resulted in *lux* bioluminescence albeit with altered kinetics (**Supplementary figure 5c and d)**. The suppression of signal observed in these experiments mirrored their impact in the cell-killing plate assays (**Supplementary figure 4**). Combined, these experiments imply that fluorophore labelling of ColE9 K469C affects the kinetics of colicin-mediated killing and that the identity of the fluorophore is a significant factor. The molecular basis of this effect is explored further below. We conclude that the residual toxicity of fluorescently-labelled ColE9 reflects a reduced capacity to transport into *E. coli* rather than the activity of unlabelled molecules.

Since fluorescently labelled ColE9 can be imported into *E. coli* we exploited this approach to probe the PMF dependence of import using fluorescence microscopy. For these experiments, ColE9 K469C^AF647^ was used because it circumvented cell auto-fluorescence issues associated with using AF488 and had lower impact on colicin-mediated killing than AF568. An active site mutation (H551A hereafter denoted as *; **Supplementary figure 3a**) was also introduced in the ColE9 K469C background to avoid killing cells during long time-courses. Following the labelling of cellular DNA with Hoechst stain, ColE9* K469C^AF647^ was added to *E. coli* JM83 cultures, the excess fluorophore removed by repeated wash steps and cells imaged on agar pads by widefield fluorescence microscopy (see Materials and Methods for details). In contrast to whole cell labelling by Hoechst, ColE9* K469C^AF647^ predominantly labelled the cell periphery (**Figure 5a and b**). To ensure labelling was specific for the ColE9 receptor BtuB, the experiment was repeated using a *btuB* deletion mutant, which abolished labelling by ColE9* K469C^AF647^ (**Figure 5c**). To determine if ColE9* K469C^AF647^ was surface exposed, labelled cells were treated with trypsin prior to imaging. Following this treatment, significant fluorescence remained associated with the cells suggesting these were imported molecules protected from proteolysis (**Figure 5a-c**). When cells were pre-treated with the protonophore CCCP, all ColE9* K469C^AF647^ fluorescence was lost from cells when trypsin was added (**Figure 5a-c**), with similar results observed for cells labelled with ColE9* K469C carrying AF488 or AF568 fluorophores (**Supplementary figure 6a and b**). Consistent with a role for PMF in ColE9* K469C^AF647^ import, pre-treatment of cells with ColB, a Ton-dependent pore-forming colicin that depolarises the inner membrane, also blocked uptake of ColE9* K469C^AF647^ across the OM similar to the effects of CCCP (**Supplementary figure 6c and d)**. As a final test for the involvement of the PMF in ColE9* K469C^AF647^ import, nigericin and valinomycin, which inhibit the proton gradient (ΔpH) and electrical potential (ΔΨ) components, respectively, were added to cells prior to labelling (**Supplementary figure 7)**. Of these, only valinomycin blocked the uptake of the colicin from which it can be inferred that it is the electrical potential of the PMF that drives ColE9 translocation across the OM. We conclude that while formation of the OM translocon complex at the cell surface is passive, import of ColE9 across the membrane is an energised process that requires the electrical potential of the PMF.

**Figure 5.**
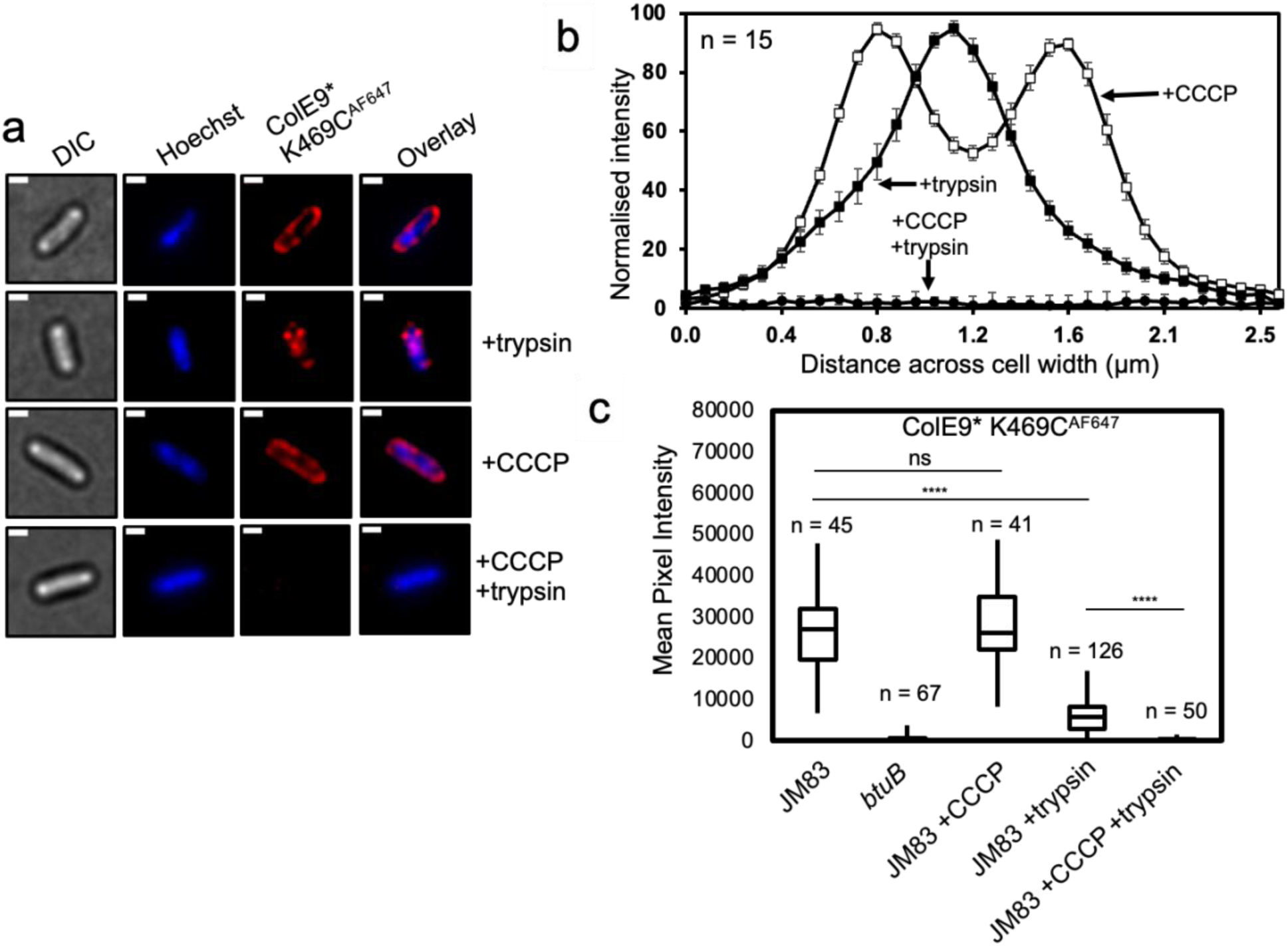
ColE9* K469C^AF647^ translocation across the outer membrane is PMF dependent. ColE9*, denotes ColE9 H551A, an active site mutation that inactivates the toxin. **a,** Widefield fluorescence microscopy images of *E. coli* JM83 cells labelled with ColE9* K469C^AF647^ (1.5 µM) and Hoechst DNA stain (20 µM) for 30 min at 37 °C with or without trypsin treatment with or without prior treatment with CCCP. Each panel show the same cell in DIC (*grey*), Hoechst DNA stain (*blue*) and ColE9* K469C^AF647^ fluorescence (*red*). Overlays of Hoechst and ColE9* K469C^AF647^ fluorescence are also shown. ColE9* K469C^AF647^ remains bound to the OM in the presence of CCCP but this signal is lost on treatment with trypsin. Trypsin treatment in the absence of CCCP yields some cell-associated ColE9* K469C^AF647^ fluorescence that likely represents internalised molecules. Scale bar, 1 µm. **b,** Fluorescence intensity across *E. coli* JM83 cell widths for each condition measured in ImageJ, n = 15 cells per condition. CCCP-treated cells showed peripheral ColE9* K469C^AF647^ fluorescence consistent with the colicin being bound at the OM. Trypsin treatment of cells in the absence of CCCP results in loss of this peripheral fluorescence but the presence of mid-cell signal consistent with internalisation. Treatment with CCCP and trypsin removed all ColE9* K469C^AF647^ fluorescence from cells, as in a. Mann-Whitney U-test performed between the +CCCP and the +trypsin data sets between cell distances of 0.8-1.6 μm, gave a value of U = 0.04, showing that there is a significant difference between these two data sets. Error bars represent % SEM. **c,** Box and whisker plots showing mean pixel intensities for ColE9* K469C^AF647^ fluorescence per cell measured for the indicated cells and condition used, whiskers represent minimum and maximum mean pixel intensity, box shows 1^st^ and 3^rd^ quartile with the median shown as a line. From left-to-right: *E. coli* JM83 cells; *E. coli btuB* deletion strain showing loss of all ColE9* K469C^AF647^ cell-associated fluorescence; *E. coli* JM83 cells in the presence of CCCP; *E. coli* JM83 cells following trypsin treatment showing significant ColE9* K469C^AF647^ fluorescence remains associated with cells indicative of import; *E. coli* JM83 cells treated with CCCP and trypsin showing the complete loss of internalised ColE9* K469C^AF647^ fluorescence. *n*, number of cells used, typically from 2-4 biological replicates. ****, indicates a P value below 0.0001 in a one-way ANOVA multiple comparisons Kruskal-Wallis test, ns indicates no significant difference.

### ColE9 translocates through the pore of a single OmpF subunit

The structural changes evident within the ColE9 translocon on displacement of BtuB, whereby the T-domain becomes repositioned immediately above the same OmpF subunit (subunit 2) as TolB in the periplasm, are consistent with OmpF being the port of entry. Indeed, OmpF eyelet mutations are known to confer resistance to Tol-dependent colicins (Jeanteur et al., 1994). The absolute requirement for the electrical potential of the PMF to import ColE9 across the OM further supports this contention since this could provide the cellular energy needed to unfold the toxin for passage through the porin. We hypothesised that if ColE9 passes directly through the eyelet then the physicochemical properties of both the colicin and the eyelet would be important factors in transport. A clue that this might be the case came from the experiments showing that three fluorophores (AF488, AF568, AF647) differentially affected the ability of ColE9 to induce the SOS response in *E. coli* DPD1718 cells (**Supplementary figure 5**); the fluorophores have molecular weights (720-1250 Da) significantly greater than are permissive in porin solute diffusion assays (Vergalli et al., 2020) yet did not impact colicin activity similarly. A complicating factor in interpreting these experiments however is the presence of both OmpF and OmpC in the OM of typical *E. coli* K-12 strains, the ratios of which change depending on the osmolarity, pH and temperature of the growth medium (Pratt et al., 1996). The two porins share 60% sequence identity, have similar trimer structures and eyelet dimensions and can form mixed trimers in the OM (Gehring and Nikaido, 1989). Where they differ however is the degree of negative charge at the eyelet; OmpC is significantly more electronegative than OmpF (**Figure 6a**). We therefore assessed the impact of fluorophore labelling on the ability of ColE9 K469C to kill *E. coli* strains expressing either OmpF or OmpC in the OM. Wild-type, unlabelled ColE9 killed both strains equally well in plate-killing assays (**Figure 6b)**. Fluorophore labelling had a modest impact on OmpF-expressing cells whereas cytotoxicity was largely abolished in OmpC-expressing cells (**Figure 6b**), especially for AF568 and AF647-labelled ColE9. These fluorophores carry -2 and -3 charges, respectively, and are the largest of the three fluorophores used (880 and 1250 Da, respectively, compared to 720 Da for AF488). Lastly, we investigated the ability of AF647-labelled ColE9 to translocate into *ompF-* or *ompC*- expressing cells using wide-field fluorescence microscopy with or without trypsin treatment. Here again, the presence of the fluorophore had a much greater impact on the ability of the colicin to enter *ompC*-expressing cells compared to those expressing *ompF* (**Figure 6c and d**). We conclude that the slightly narrower pore and greater negative charge of the OmpC eyelet act as significant impediments to ColE9 import when the colicin has a large (>600 Da), negatively charged fluorophore covalently attached to its C-terminal DNase domain, an attachment that has no effect on nuclease activity (**Supplementary figure 3**). Since ColE9 import is dependent on the PMF and delivery of the PMF occurs via the N-terminal regions of the colicin, these results can only be explained if the entire colicin passes through the eyelet of the porin carrying the fluorophore with it.

**Figure 6.**
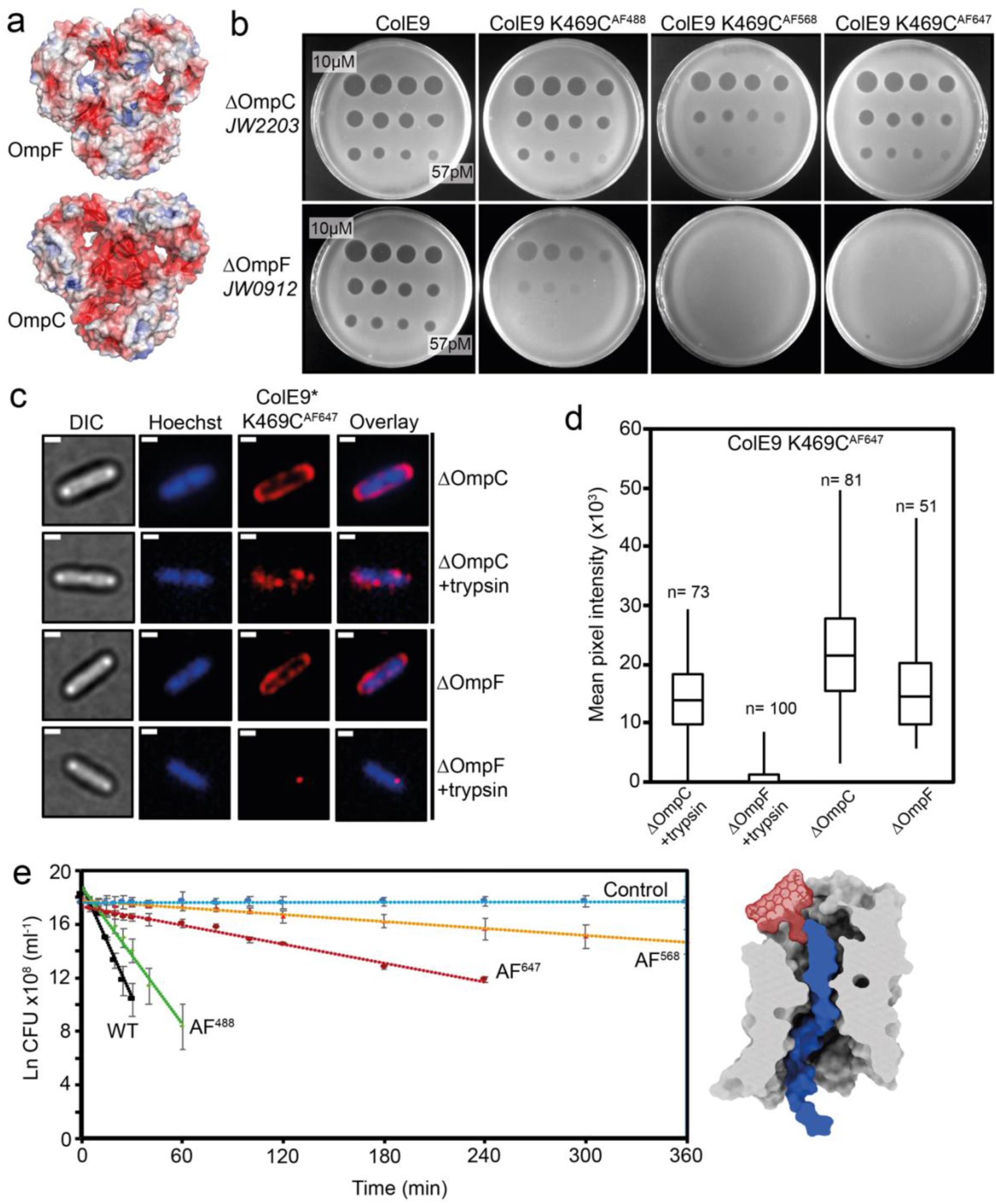
Labelling ColE9 with fluorophores impedes its translocation through the pores of OmpC. **a**, Cut-through molecular surface images showing the distribution of charged residues (acidic, *red*; basic, *blue*) at the eyelets of trimeric OmpF (PDB ID 3K19) and OmpC (PDB ID 2J1N). The eyelet of OmpC is significantly more electronegative than that of OmpF. Electrostatic surfaces were calculated using the APBS plugin within Pymol. **b**, Overnight plate assays comparing the cytotoxic activity of wild-type ColE9 with ColE9 K469C labelled with AF488, AF568 or AF647 against a lawn of *E. coli* with either OmpF (JW2203) or OmpC (JW0912) in the outer membrane. Each plate was spotted with a serial dilution of the colicin (10 µM - 57 pM). Wild-type ColE9 was equally active against both ΔOmpF and ΔOmpC cells, whereas labelling ColE9 K469C with fluorophores reduced (AF488) or abolished (AF568, AF647) colicin activity in ΔOmpF cells, whilst activity against ΔOmpC cells remained comparable to that of wild-type ColE9. **c,** Widefield fluorescence microscopy images of *E. coli* JW2203 (*ompF*-expressing) and JW0912 (*ompC*-expressing) cells labelled with ColE9* K469C^AF647^ (1.5 µM) and Hoechst stain (20 µM) for 30 min at 37 °C with or without trypsin treatment. Each panel shows the same cell in, DIC (*grey*), Hoechst DNA stain (*blue*) and fluorescence of ColE9* K469C^AF647^ (*red*). Overlays of Hoechst and ColE9* K469C^AF647^ fluorescence are also shown. Data show that following trypsin-treatment significant ColE9* K469C^AF647^ fluorescence remains associated with *ompF*-expressing *E. coli* whereas little or no fluorescence remains associated with *ompC*-expressing *E. coli*. Scale bar, 1 µm. **d**, Box and whisker plots of *E. coli* JW2203 and *E. coli* JW0912 cells labelled with ColE9* K469C^AF647^ with and without trypsin treatment: whiskers represent minimum and maximum mean pixel intensity, box shows 1^st^ and 3^rd^ quartile with the median shown as a line. Microscopy data were collected as in c. The mean pixel intensity of ColE9* K469C^AF647^ per cell was measured for each cell condition. *n*, number of cells, typically from 3 or 4 biological replicates. Data show that ColE9* K469C^AF647^ translocates across the OM through OmpF and that this is significantly impeded when *E. coli* has OmpC in the outer membrane. **e**, Shows first-order cell death kinetics for wild-type ColE9 or ColE9 K469C labelled with different AF dyes (AF488, AF568, AF647) against *E. coli* JW2203 *ompF*-expressing cells. Cultures were incubated with 80 nM toxin at pH 7.5 and 37 °C and at various time points the reaction stopped using trypsin at 37 °C for 30 min, cells plated out and CFUs recorded. All time courses were conducted in the presence of chloramphenicol (20 µg/ml) to reversibly block cell division, which would otherwise compete with cell-killing. Control data shown are for cells with no ColE9 added but where growth was inhibited by the presence of chloramphenicol. Cell-killing half-lives, obtained from the fitted first-order plots shown (error bars from two biological replicates), were as follows: wild-type ColE9, 2.5 min; ColE9 K469C^AF488^, 4.1 min; ColE9 K469C^AF647^, 29.6 min; ColE9 K469C^AF568^, 78.8 min. The data show that the chemical nature of the fluorophore in the C-terminal DNase domain of ColE9 has a dramatic effect on the cell-killing kinetics of the colicin. Schematic surface representation in which AF568 (model Generated in coot), the fluorophore with the biggest impact on OmpF-mediated killing, has been manually grafted onto the structure of ColE9 OBS2 bound within subunit 2 of OmpF. The slow cell-death kinetics of ColE9 K469C^AF568^ likely reflects the time taken to pass this bulky molecule through the narrow eyelet of the porin.

The differential effects of fluorophores on ColE9 toxicity and import in cells harbouring OmpC or OmpF raised the question of why, when the dimensions of porin eyelets are similar, bulky fluorophores pass apparently unhindered through OmpF? We hypothesized that the long time-course (overnight) of the cell-killing assays might mask the import effects of bulky fluorophores on *E. coli* cells expressing *ompF*. To determine if more subtle effects were evident on shorter timescales the kinetics of killing *E. coli* JW2203 cells, which only express *ompF*, were determined after addition of ColE9 K469C labelled with AF488, AF568 or AF647 (**Figure 6e**). The kinetics of cell death were monitored by measuring colony forming units (CFUs) as a function of time, following the addition of trypsin to degrade extracellular colicin. Chloramphenicol was used throughout these experiments as a bacteriostatic agent to stop cell division, which would otherwise complicate interpretation of cell-death kinetics. The data in **Figure 6e** demonstrate that the ColE9-mediated killing of *ompF*-expressing cells showed a first-order kinetic profile regardless of which fluorophore is attached. The fluorophores nevertheless had a profound impact on the half-life of this process; the half-life for cell-death by unlabelled ColE9 was ∼2.5 min, at 37°C, which increased almost two-fold for ColE9 K469C^AF488^, ∼10-fold for ColE9 K469C^AF647^ and ∼30-fold for ColE9 K469C^AF568^. The relative order of fluorophore impact on the killing half-life (AF568>AF647>AF488) was the same as the order observed in the plate killing assay with *E. coli* JM83 cells (**Supplementary figure 4**) and the induction of the SOS response in *E. coli* DPD1718 cells (**Supplementary figure 5**), where both strains express *ompF* and *ompC*. In conclusion, the kinetic profiles shown in **Figure 6e** reflect the rate-limited, energised transport of ColE9 molecules through the lumen of a single OmpF subunit following assembly of the OM translocon, which is a concept that is further explored in the Discussion.

## Discussion

### Model for the force-dependent import of ColE9 through OmpF

Combining cryo-EM structure determination of ColE9 translocon complexes with fluorescence microscopy data, and in conjunction with past biochemical, biophysical and structural data on this and related colicins, it is finally possible to unravel the complex interplay of passive and active processes at play in the entry mechanism of Tol-dependent colicins (**Figure 7**).

**Figure 7.**
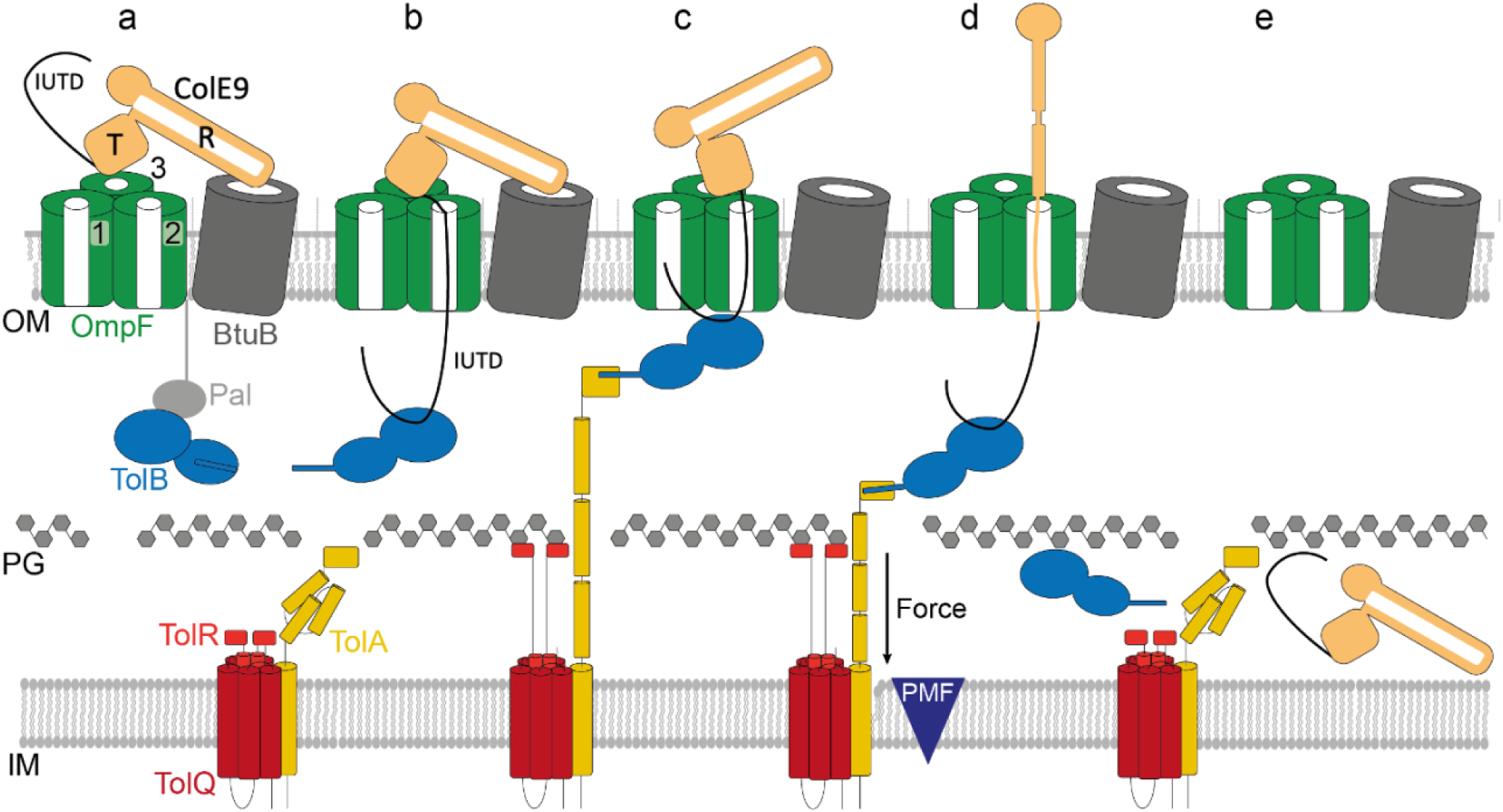
Model of ColE9 translocon assembly and energised OM transport (see text for details). **a,** ColE9 R-domain binds BtuB with high affinity, positioning the T-domain and IUTD above a neighbouring OmpF trimer. **b**, The ColE9 IUTD translocates through subunit 2 of OmpF to deposit the TBE in the periplasm and capture TolB at the expense of the OM lipoprotein Pal. The TBE allosterically promotes displacement of TolB’s N-terminus which constitutes the TolA binding site. We propose this complex equates to the full translocon cryo-EM structure. **c**, ColE9 OBS1 binds subunit 1 of OmpF from the periplasm. Threading through two of OmpFs three subunits drives dissociation of the ColE9 R-domain-BtuB complex and reorients the T-domain above subunit 2 below which TolB is positioned in the periplasm. The docking of OBS1 also results in TolB moving closer to the opening of subunit 2. This complex, which we propose equates to the partial translocon cryo-EM structure, is now primed for contact with TolA in the inner membrane. TolA extension through the periplasm is coupled to the PMF via its stator proteins, TolQ and TolR. **d**, Retraction of TolA (the molecular mechanism of which remains to be established) provides the driving force for pulling ColE9 bound to TolB into the periplasm through subunit 2 of OmpF, accompanied by unfolding of its constituent domains. It is at this point the immunity protein Im9 (not shown) would be displaced at the cell surface. **e**, The TolA-TolB complex is thought to retract through the cell wall, which would bring ColE9 close to the cytoplasmic membrane. The toxin likely refolds prior to transport across the cytoplasmic membrane, which involves the AAA^+^ATPase/protease FtsH (not shown) (Walker et al., 2007).

Past work has shown that E group colicins assemble their OM translocons by first binding with nM affinity to BtuB at a 45° angle relative to the plane of the membrane from where the N-terminal IUTD captures a neighbouring OmpF (**Figure 7a**) (Housden et al., 2005; Kurisu et al., 2003). The cryo-EM structure of the intact ColE9 translocon reveals that the colicin also has to overcome a 35° tilt of BtuB relative to OmpF, which it does by having an R-domain with long enough reach. From this overhanging position, the IUTD associates with the pore of an OmpF subunit (designated subunit 2 in our structure), most likely due to its high local concentration and electrostatic attraction between the pore which is mildly cationic selective and the N-terminus of the colicin. The disordered polypeptide chain then reptates through subunit 2 of OmpF to deliver the TBE to the periplasm, capturing TolB at the expense of its endogenous binding partner, the OM lipoprotein Pal (**Figure 7b**) (Bonsor et al., 2007; Bonsor et al., 2009). ColE9 mimics the interactions of Pal, unlike Pal however the colicin induces TolB to expose its N-terminus which houses the TolA binding site. Concomitant with (or following) binding to TolB, the ColE9 OBS1 sequence inserts into subunit 1 of OmpF and OBS2 becomes stably bound within subunit 2. The R-domain of ColE9 dissociates from BtuB thereby accommodating OmpF threading and removing an architectural constraint on the repositioning of its T-domain, from a central position above OmpF to one that is now directly above subunit 2 (**Figure 7c**). OmpF threading is also accompanied by rotation of the ColE9 TBE-TolB complex in the periplasm and formation of additional stabilising interactions between the colicin and the porin, all of which position the ColE9 TBE-TolB complex close to the periplasmic entrance of OmpF subunit 2. It is at this point that optimal engagement between PMF-linked TolQ-TolR-TolA in the inner membrane and the ColE9 TBE-TolB complex likely occurs (**Figure 7c**). The physiological role of the TolQ-TolR-TolA assembly is to drive dissociation of TolB-Pal complexes at the OM, releasing Pal to bind the cell wall (Szczepaniak et al., 2020). As a consequence of this PMF- driven cycle, TolB is thought to be pulled through the cell wall by TolA. Hence, by virtue of its association with TolB, ColE9 is dragged unfolded into the periplasm through subunit 2 of OmpF, its C-terminal DNase domain the last to enter the cell (**Figure 7d, e**). Previous colicin translocation models have suggested Tol-dependent bacteriocins remain associated with their OM components while killing the cell (Benedetti et al., 1992; Duche, 2007). Our data suggest this is unlikely to be the case. The entire ColE9 molecule enters the periplasm, driven by interactions at its N-terminus with the PMF-driven Tol-Pal system. Subsequent transport of the nuclease domain across the inner membrane requires the AAA^+^ATPase/protease FtsH. FtsH also proteolytically processes the domain as it enters the cytoplasm, although the underlying mechanism is unknown (Chauleau et al., 2011; Walker et al., 2007).

Forced unfolding of ColE9 during OM transport is consistent with several lines of evidence. First, our cryo-EM data. The structural changes that accompany ColE9 translocon formation, in which the T-domain and ColE9 TBE-TolB complex become aligned over the same OmpF subunit, suggest they are the means by which the bacteriocin maximises its access to PMF-mediated force transduction through OmpF. Second, internal disulphide bonds within the coiled-coil domain block ColE9 entry, which can be explained by the crosslinks blocking the unfolding necessary to allow entry through OmpF (Penfold et al., 2004). Third, *in vivo* experiments have shown that release of Im9 (which forms a fM complex with the E9 DNase) at the cell surface is dependent on the electrical potential of the PMF (Vankemmelbeke et al., 2013), which is also the case for the entry of fluorescently-labelled ColE9 in our experiments. These observations are consistent with the release of Im9 accompanying the force-dependent unfolding and import of the colicin DNase through OmpF. Fourth, *in vitro* atomic force microscopy measurements have demonstrated that when forces equivalent to those attainable by the PMF are applied to the isolated ColE9 DNase-Im9 complex the DNase domain melts, releasing Im9 but only when the force is applied at the N-terminus of the domain (Farrance et al., 2013). Viewed in the context of a translocating ColE9 molecule at the OM, the force exerted by the PMF would similarly be applied at the N-terminus of the toxin, with the surface of OmpF acting as the barrier against which the toxin is unfolded and Im9 displaced.

We conclude that the transport mechanism uncovered for ColE9 is likely to be the basis for the import of most Tol-dependent bacteriocins, the majority of which require porins. This will certainly be the case for E group nuclease colicins (E2-E9) that use BtuB as their receptor, and share high sequence identity with ColE9, but deliver rRNases or tRNases as well as DNases into the cell. This is also likely to be the case for pore-forming bacteriocins such as ColA, ColK and ColN, which use porins as translocators but different OM receptors (Kleanthous, 2010).

### Ton- and Tol-dependent bacteriocins exploit analogous PMF-dependent mechanisms to cross the outer membrane through protein pores

The basic import principles we have uncovered for ColE9 highlights similarities with those recently described for the Ton-dependent bacteriocins pyocins S2 and S5, which target *Pseudomonas aeruginosa*. Both S2 and S5 associate with the common polysaccharide antigen on the cell surface and then exploit different TBDTs, FpvAI and FptA, respectively, as translocators to the periplasm (Behrens et al., 2020; White et al., 2017). Translocation of these bacteriocins involves two energised events. The first displaces the plug domain that occludes the central pore of the TBDT by a TonB-dependent mechanism, akin to that accompanying ligand uptake through the transporter. The second involves transport of the unfolded toxin through the now-open TBDT pore following delivery of the toxin’s N-terminal TonB box to the periplasm. The mechanism we describe for Tol-dependent bacteriocins is analogous to the second of these steps. The similarity is underscored by the way in which force is exerted by TonB and TolA on their endogenous OM protein targets. The C-terminal domains of both proteins form structurally related, parallel β-sheets with their partners involving β-strand augmentation; TonB with the N-termini of TBDTs, TolA with the N-terminus of TolB (Pawelek et al., 2006; Szczepaniak et al., 2020). Such complexes are resistant to mechanical deformation (Chen et al., 2015). Hence, when the PMF-coupled stators ExbB-ExbD and TolQ-TolR, respectively, exert force on them instead of dissociating they remain bound, descending towards the inner membrane as TonB and TolA retract (Hickman et al., 2017). By hijacking these systems, both Ton- and Tol-dependent bacteriocins use the PMF to import themselves across the OM in unfolded form through either pre-existing pores (porins) or pores opened transiently by the PMF (TBDTs).

### Rate-limited transport of ColE9 through porins can be used to import large chemical entities into *E. coli*

The single-hit kinetic profiles observed for colicin-mediated killing of *E. coli* have been known since the 1960s (Reeves, 1965). However, the basis of this characteristic phenomenon has remained elusive. Our data reveal that this kinetic behaviour reflects the rate-limited transport of the colicin through the OM. The cell-killing half-life for ColE9 targeting *E. coli* cells with OmpF as the only porin in the OM (∼2.5 min at 37 °C and pH 7.5) changes significantly when AlexaFluor dyes are attached to the DNase domain, by up to 30- fold in the case of AF568. We suggest that much of this additional time reflects repeated attempts by the Tol-Pal system, in conjunction with the PMF, to bring this single, bulky, negatively-charged adduct through the narrow eyelet of OmpF subunit 2 (**Figure 6e**). The task of bringing this adduct through OmpC becomes even greater because of the eyelet’s greater negative charge.

Glycopeptide antibiotics like vancomycin do not kill Gram-negative bacteria because they cannot partition into the OM and are too large to diffuse through porins. This impermeability problem is a major factor in multidrug resistance amongst Gram-negative pathogens. One approach being developed to circumvent these issues is to conjugate such complex antibiotics to siderophores (so-called Trojan horse antibiotics) so that they can be actively transported through specific TBDTs, energised by the PMF-linked Ton-system (Klebba et al., 2021). Our data show that large, bulky, organic fluorophores that approach the size and complexity of vancomycin can be actively transported through the porins of *E. coli* by piggy-backing on a colicin. Colicins might therefore represent a new vehicle for the delivery of antibiotics that cannot otherwise penetrate the OM of Gram-negative bacteria.

## Acknowledgements

We are indebted to Patrice Rassam (Strasbourg) and Patricks Inns (Oxford) for help and advice regarding microscopy experiments, David Staunton (Molecular Biophysics Suite, Oxford) for help with biophysical analysis of purified and fluorescently-labelled proteins, and Jeremy Keown and Loic Carrique for assistance with computational analysis of cryo-EM data. Cryo-EM data for this investigation were collected at the ISMB EM facility at Birkbeck College, University of London with financial support from Wellcome Trust (202679/Z/16/Z and 206166/Z/17/Z). The work was funded by the European Research Council (Advanced grant 742555; OMPorg) and the Wellcome Trust (201505/Z/16/Z). MLRF was supported by the BBSRC Oxford Interdisciplinary Bioscience DTP. We are also indebted to Nadia Halidi and the Micron Advanced Bioimaging Unit (Wellcome Strategic Awards 091911/B/10/Z and 107457/Z/15/Z) for their support & assistance in this work. We thank the Oxford Biomedical Research Computing (BMRC) facility, a joint development between the Wellcome Centre for Human Genetics and the Big Data Institute (supported by Health Data Research UK and the NIHR Oxford Biomedical Research Centre).

## Author contributions

N.G.H, M.N.W., M.L.R.F, N.L. and C.K. designed research; R.K., E.E. and B.C. provided materials; M.L.R.F., M.N.W., and N.L. performed research; M.L.R.F and M.N.W. analysed data; C.K. and M.N.W. wrote the paper.

## Data Availability Statement

The data supporting the findings of the study are available in the article, available upon request from the corresponding author or submitted to the protein structure data bank (PDB).

## Materials and Methods

### Plasmids, bacterial strains and media

Whole plasmid site-directed mutagenesis was used to introduce a solvent accessible cysteine K469C in the cytotoxic domain of ColE9.Im9His6, cloned into pET21a to give pMLF07. pMLF07 was used to introduce H551A within the cytotoxic domain to give pMLF08, a cytotoxic inactive construct. Plasmid pEE01 was used to express ColB-His6 in pACYCduet-1. Plasmid pBC1 expressing ColE9 DNase K469C.Im9His6, containing ColE9 DNase domain with K469C single cysteine, in pET21d, was generated from pRJ353. Plasmid pREN151 was amplified from a pCS4 derivative and inserted between NdeI and XhoI sites of pET21a to enable expression and purification with C-terminal His6 tag. All *E. coli* strains and plasmids used in this study are listed in supplementary tables 2 and 3. Strains were routinely grown at 37 °C in either Lysogeny broth (LB) (Miller 1972) or M9- minimal media or plated on LB-agar. Supplemented M9-minimal media contained 47.78 mM Na_2_HPO_4_, 22 mM KH_2_PO_4_, 8.56 mM NaCl, 2 mM MgSO_4_, 0.1 mM CaCl_2_, 18.7 mM NH_4_Cl, 0.4 % (w/v) D-glucose, 0.05% (w/v) casamino acids and 0.0002 % (w/v) thiamine.

### Protein purification

ColE9 and TolB variants were expressed and purified as described previously (Garinot-Schneider et al., 1997; Garinot-Schneider et al., 1996b; Housden et al., 2013; Loftus et al., 2006). Plasmids encoding protein of interest were expressed in *E. coli* BL21 (DE3) cells grown in shaking LB flasks at 37 °C (supplemented with 100µg/ml ampicillin) to an OD_600_ of 0.7 and protein expression induced with 1 mM isopropyl-β-D-1-thiogalactopyranoside (IPTG). Cells were grown for a further 3 h, then harvested by centrifugation, re-suspended in binding buffer (20 mM potassium phosphate pH 7, 500 mM NaCl, 5 mM imidazole) supplemented with1 mM PMSF (and 10mg lysozyme for pMLF07/08) and lysed by sonication. The soluble cell lysate obtained after centrifugation was loaded onto a 5 ml HisTrap FF HP nickel column (GE Healthcare) equilibrated with binding buffer. ColE9 K469C and ColE9 H551A K469C were eluted from column-bound His-Im9 through denaturation with guanidine hydrochloride (20 mM potassium phosphate pH 7, 500 mM NaCl, 6 M guanidine pH 7). Eluted fractions were analysed by SDS-PAGE to locate protein before overnight dialysis into SEC buffer to refold protein (20 mM potassium phosphate pH 7, 500 mM NaCl) supplemented with 10 mM DTT. For ColE9 W39A and ColE9 R-domain (pNGH96 and pRK151) were eluted using a linear imidazole gradient from 50-250 mM over 10 column volumes. Following SDS-PAGE analysis, the pooled sample was spiked with EDTA (5 mM) and dialysed overnight into SEC buffer containing 20 mM Tris/HCl pH 7.5, 150 mM NaCl. Proteins were further purified by SEC, using a Superdex 200 Hiload 26/600 column (GE Healthcare) equilibrated with SEC buffer (10 mM DTT added to buffer where ColE9 contained free cysteine residue). Protein concentrations were determined by UV absorbance at 280 nm using sequence-based extinction coefficient (Gill and von Hippel, 1989).

### Preparation of the ColE9 translocon for Cryo-EM

Intact ColE9 translocon complex was prepared as previously described (Housden et al., 2013). The purified ColE9 translocon in 20 mM MES pH 6.5 and 1% β-OG (100 µL), was diluted 10-fold into buffer containing 20 mM potassium phosphate, pH 7.9, 100 mM NaCl and 1% β-OG. Amphipol was added to protein in a ratio of 10 mg amphipol:1mg protein, and the sample was left to incubate at room temperature for 2 hr. Detergent was removed through the addition of BioBeads (Bio-Rad) at a ratio of 10 mg beads to 1 mg of detergent. After a 2 hr incubation at room temperature, the sample was injected onto a superose 6 increase 10/300 column, equilibrated in 20 mM potassium phosphate, pH 7.9, 100 mM NaCl. Fractions were analysed by SDS-PAGE to validate all components of the complex were present, before spin concentrating the sample to 2.0 mg/ml.

The quality and concentration of samples were confirmed by negative stain EM prior to cryo- EM sample preparation. C-flat grids (Protochips, USA; 1.2/1.3 300 mesh) and UltrAuFoil (Quantifoil, Germany; 1.2/1.3 300 mesh) were negatively glow discharged using PELCO Easiglow (Ted Pella, USA) and coated with graphene oxide as described previously (Wang et al., 2020). Prior to application to graphene coated quantifoil grids, translocon preparation was diluted to 0.05 mg/ml with buffer containing 20 mM potassium phosphate, pH 7.9, 100 mM NaCl. 3 µL of sample was applied to grid and vitrified in liquid ethane using a Vitrobot Mark IV (Thermo Fisher Scientific, USA) at 4°C and 94% humidity.

### Cryo-EM image collection and processing

Cryo-EM data were collected in ISMB Birkbeck EM facility using a Titan Krios microscope (Thermo Fisher Scientific, USA) operated at 300 keV and equipped with a BioQuantum energy filter (Gatan, USA). The images were collected with a post-GIF K2 Summit direct electron detector (Gatan, USA) operated in counting mode, at a nominal magnification of 130,000 corresponding to a pixel size of 1.047 Å. The dose rate was set to 4.99 e Å^2^ per second, and a total dose of 49-50 e/Å^2^ was fractionated over 50 frames. An energy slit with a 20 eV width was used during data collection. Data were collected using EPU software with a nominal defocus range -1.5 µm to -3.5 µm. Relion 3.1 software package (Scheres, 2012a, b; Zivanov et al., 2018; Zivanov et al., 2020) was used for data motion correction and dose weighting with MotionCor2 (Zheng et al., 2017). Data were then imported into cryoSPARC (Punjani et al., 2017; Punjani et al., 2020) and the contrast transfer function (CTF) estimated using GCTF (Zhang, 2016). After removing micrographs with non-vitreous ice, broken graphene oxide support or poor CTF fit, blob picking followed by particle extraction, and 2D classification were used to generate templates for template-based picking within cryoSPARC. Template-picked particles were extracted with a 280- pixel box and sorted using 2D classification. From here, the processing workflow for the two datasets diverged.

For data set one (654,865 particles), five *ab initio* classes were generated following 2D classification, and classes were refined using heterologous refinement within cryoSPARC. In two of the resulting classes, the density was consistent with a translocon complex, these were selected for further processing using non-uniform refinement, prior to import into relion3.1. After 3D refinement the full translocon map reported an overall resolution of 5.0 Å and the partial translocon map, 3.7 Å. To improve the quality of the partial translocon map the detergent shell was subtracted and the resulting particles were refined to generate a new map with a reported resolution of 3.9 Å. In parallel to 3D refinement subtracted particles were also further sorted using a 3D classification step, without alignment and sorting into four classes. All four classes were refined independently, with only one class showing map density for both ColE9 and TolB. After a 3D refinement of this class, a map was generated with a reported resolution of 3.7 Å which, following post-processing and local resolution estimation, was sharpened using either LocScale (Jakobi et al., 2017) or deepEMhancer (Sanchez-Garcia et al., 2020). Visual inspection of the sharpened maps confirmed the selection of the deepEMhancer tight map (**Supplementary figure 1**) as the final map for the partial translocon. Similar sorting and refinement procedures were applied to the full translocon particles, however no improvements in map resolution and quality were observed.

Data set two (434,671 particles after 2D classification) was processed using cryoSPARC. Three *ab initio* maps were generated and subject to heterologous refinement. Of these classes only one class showed weak density for both TolB and ColE9, at higher threshold levels, in addition to the density consistent with the OmpF trimer. This map was then further processed using non-uniform refinement. The resulting map reported a resolution of 3.2 Å, however upon visual inspection of the map it revealed poor density for TolB and ColE9 relative to the partial translocon map from dataset one, and also had non-isotropic density within OmpF, therefore no further processing was carried out.

After independent processing of both datasets, the particles were combined to give 1,089,536 particles which were sorted using a second 2D classification step. Classes were divided into three groups, full translocon, partial translocon, and junk. The particles in the full and partial translocon classes were independently processed within cryoSPARC. Two maps of the full translocon were generated from *ab initio* reconstruction. One map had a more intact structure with density for all components and was therefore refined further using non-uniform refinement, to give a map with a reported resolution of 4.0 Å. Visual inspection of the map showed that only the OmpF portion of the map extended to 4.0 Å with the majority of the remaining map having poorly resolved secondary structure and weak density associated with ColE9 and TolB. To improve the quality of the map, the detergent shell was subtracted and the map locally refined. This generated a map with a reported resolution of 4.6-16 Å, however the density of ColE9 and TolB had improved. This map was sharpened with either LocScale or deepEMhancer, with the deepEMhancer tight mask map generating the final map of the full translocon. To assess the degree of conformational heterogeneity, 3D variability analysis was implemented using particles input into the final map. From the analysis five maps were generated, which showed movement within TolB and ColE9. Comparison of these maps with that of the partial translocon, did not show a map whereby the conformation of ColE9 or TolB matched that seen in the partial translocon map, suggesting that the partial translocon and full translocon were discrete states.

After combining datasets, particles assigned to the partial translocon class were processed using a similar method to that applied to the full translocon above. Four classes were generated from *ab initio* reconstruction and heterologous refinement. Of these four classes only one showed density for both ColE9 and TolB, this class was processed further using non-uniform refinement, local refinement, detergent shell subtraction, and a final local refinement step, to generate a map with a reported resolution of 3.6 Å. This map was input into 3D variability analysis generating five maps to assess the degree of conformational heterogeneity. Once again density for ColE9 consistent with the full translocon map was absent.

To build a partial translocon model OmpF (PDB ID 3K19), was docked into the partial translocon map from relion using chimera (Pettersen et al., 2004) and rigid-body refined using Phenix (Afonine et al., 2018a; Afonine et al., 2018b; Liebschner et al., 2019). ColE9 residues 2-75 were manually built into the remaining map density present within the OmpF pores within coot (Emsley and Cowtan, 2004) and several cycles of rigid-body and local refinement was carried out using Phenix. TolB-ColE9 32-47 (PDB ID 2IVZ) was then docked into map using chimera and rigid body refined using phenix. Due to the weak density in the TolB portion of the map, the ColE9 peptide was not clearly resolved, however after rigid-body fitting the terminal residues (32 and 47) were proximal to the ColE9 chain built prior to TolB docking. Therefore, residues 32-47 of ColE9 bound in the TolB crystal structure were substituted for *ab initio* built sequence and the whole complex (OmpF, TolB and ColE9 residues 2-75) refined using phenix. Only rigid body refinement was applied to the TolB portion of the map due to the lower resolution in this region of the map. The ColE9 T-domain residues 85-314 (PDB ID 5EW5) was docked into the map using chimera, merged with remaining parts of the translocon, and rigid body refined in phenix to give the final partial translocon model. Due to the weak density following on from ColE9 residue 75 the disordered N-terminal domain could not be reliably built past this point to connect residues 75 and 85.

For the full translocon model, OmpF and ColE9 residues 2-75 from the partial translocon structure were docked into the final full translocon map within chimera. Additionally, BtuB bound to the R-domain of ColE3 (PDB ID 2YSU) was also docked into the map, prior to running refinement within Phenix. Only rigid body refinements were run for this model due to the lower resolution of the map. The crystal structure of ColE9 (PDB ID 5EW5) was docked into the map, through alignment to the R-domain of ColE3 within chimera. This created an anchor point for the R-domain that prevented steric clashes with BtuB. The TolB-ColE9 32-47 crystal structure was once again docked into the map and merged with ColE9 2-75, and the whole complex rigid-body refined in phenix, to generate the final model.

Map to model parameters from Phenix, model validation and map processing workflow are summarised in **Supplementary Figures 1-3** and **Supplementary Table 1**. Structure representation figures were made using chimeraX (Goddard et al., 2018; Pettersen et al., 2021) and pymol.

### Vitamin B_12_ competition assay

The assay was carried out as described previously (Penfold et al., 2000). In brief, *E. coli* 133/3 cells grown were overnight in 5 ml minimal media (1x M9 salts, 100 µg/ml L-arginine, 1 mM MgSO_4_, 0.2% glucose) and supplemented with vitamin B_12_ (1 nM), prior to dilution 1:100 in 50 ml of minimal media containing vitamin B_12_ (1 nM) and ColE9 R-domain or ColE9 W39A (40 nM). Cells were maintained at 37 °C with shaking for a total time of 6 hours. Samples (1 ml) were removed every 30 minutes and optical density at 600 nm (OD_600_) recorded. Average OD_600_ values and standard deviation across triplicate measurements are reported.

### Colicin cytotoxicity assay

Colicin cytotoxicity was assayed by plate-based growth inhibition assays that were carried out as described previously (Atanaskovic et al., 2020; Jansen et al., 2020) An overnight culture of *E. coli* was used to inoculate LB (5-10 ml, containing appropriate antibiotics) and grown at 37 °C until reaching an OD_600_ ∼0.4. 200 µL of culture was added to 5 ml of molten soft LB-agar (0.75 % (w/v)) at 45 °C, mixed by inversion and poured over LB-agar plates (supplemented with appropriate antibiotics) and allowed to cool and dry. Serial dilutions of colicin were spotted onto the plate (2 µl) and incubated at 37 °C overnight. Colicin induced cell killing was indicated as zones of clearance in the bacterial lawn.

### Fluorophore labelling of proteins

Single cysteine mutants of ColE9 or the ColE9 DNase domain were desalted on a 5 ml HiTrap column (GE Healthcare) equilibrated with binding buffer (20 mM potassium phosphate pH 7, 500 mM NaCl) to remove DTT. The protein was diluted to 50 µM and labelled with a 5:1 molar excess of fluorophore:protein with either AlexaFluor 488 C_5_- Maleimide (Invitrogen) (AF^488^), AlexaFluor 568 C_5_-Maleimide (Invitrogen) (AF^568^), or AlexaFluor 647 C_2_-Maleimide (Invitrogen) (AF^647^). Maleimide dyes were re-suspended in DMSO and stored as a 10 mM stock. Labelling was allowed to proceed for 1-2 hours at room temperature in the dark, and then quenched using 2 mM DTT for 15 min, before overnight dialysis at 4°C (20 mM potassium phosphate pH 7, 500 mM NaCl) to remove excess dye (Spectra Por 10K MWCO). Labelled protein was applied to a Superdex 200 10/300 GL column (GE Healthcare) equilibrated with 20 mM potassium phosphate pH 7, 500 mM NaCl, to remove residual dye. Protein-containing fractions were identified by SDS-PAGE. The protein concentration and labelling efficiency (typically >90%) was estimated from absorbance readings using sequence-based extinction coefficients, molecular weights, and dye correction factors (Invitrogen)

### Plasmid nicking assay

pUC19 DNA was used at 1 µg/ml in 50 mM triethanolamine (Fisher Scientific) pH 7.5 with 20 mM MgCl_2_ and 200 nM ColE9 DNase domain. 5 µl samples were taken at timepoints, the reaction quenched with EDTA (37.5 mM, pH 8.0) and products separated on an agarose gel (1.2 %) following the addition DNA loading dye (Invitrogen).

### Bioluminescence assay

*E. coli* DPD1718 cells were inoculated from an overnight culture, into 10 ml LB and grown at 37 °C until OD_600_ ∼0.4. Cells were diluted 1:2 in 100 µl with warmed LB (37 °C), into a black Nunc microwell 96 well optical bottom polystyrene plate. ColE9 was used at 0, 0.5, 1, 2, and 20 nM. The plate was loaded into a plate incubator (Freedom Evo 150 robot) at 37 °C with gentle shaking. After loading of plates into Tecan Infinite M200 Pro plate reader cell growth (600 nm) and bioluminescence (490 nm) measured every 10 minutes.

### Cell preparation for live cell imaging by fluorescence microscopy

*E. coli* cells from an overnight LB culture were used to inoculate 10 ml of supplemented M9 minimal media (with the appropriate antibiotics) and grown at 37 °C until they reached OD_600_ of 0.4. Cells (1.6 ml) were pelleted by centrifugation (7000 g for 3 mins at 17 °C), and re-suspended in 200 µl M9 containing 1.5 µM of fluorescently-labelled colicin and 20 µM Hoechst 33342 (Thermofisher). Cells were incubated for 30 minutes at room temperature with rotary inversion in the dark, then pelleted by centrifugation and re-suspended in 400 µl supplemented M9 minimal media. Pelleting followed by resuspension was repeated twice more to remove unbound label. Finally, pelleted cells were re-suspended in a small volume (10-100 µl) of M9 and 10 µl loaded onto 1% (w/v) M9 minimal media agarose pad using a 1.5 x 1.6 cm Gene Frame matrix, solidified on a 1.2-1.5 mm thick microscope slide (Thermo Fisher).

For CCCP experiments, pelleted cells were re-suspended in 400 μl of 100 μM CCCP (Sigma) and incubated for 5 minutes at room temperature prior to the addition of ColE9. CCCP remained present in media throughout wash steps and in the agarose pad. For nigericin and valinomycin experiments, cells were permeabilised using 1 mM EDTA pH 8.0 for 5 minutes before treatment with 5 μM nigericin (Sigma) or 5 μM valinomycin (Sigma) in the presence of 150 mM KCl, and incubated for 5 minutes at room temperature, to inhibit the proton concentration gradient (ΔpH) or the electrical potential (ΔΨ), respectively, before addition of colicin. Nigericin and valinomycin remained present throughout the washes and in the agarose pad.

For trypsin experiments, labelled and washed cells were re-suspended in 200 µl M9 media containing trypsin (Sigma) to achieve a 2:1 colicin:trypsin ratio. Cells were incubated with trypsin for 1 h at room temperature with rotary inversion before washing three times in M9.

### Widefield microscopy data acquisition and analysis

Live cells were imaged at room temperature using a 60x oil-immersion objective (Oil UPlanSApo) (NA 1.42) on an OMX V3 BLAZE Optical microscope with Deltavision OMX software. The 405 nm (DAPI filter), 488 nm (FITC filter), 593 nm (615/24 filter) or 642 nm (Cy5 filter) laser lines were used. Lasers were used at 10-31.5 % laser intensities and an exposure time of 30 ms. Differential interference contrast (DIC) was used to image the cell. The fluorescence channels were used to acquire single images, recorded at a frame rate of 30 ms/frame.

Widefield data in the fluorescence channels (405 nm, 488 nm, 593 nm and 642 nm) were smoothed using ImageJ (Schneider et al., 2012). The data were normalised and set at a minimum of ∼100 units/pixels to remove background. Fluorescence intensity distribution across the width of cells was determined using ImageJ (Schneider et al., 2012) plot profile function. A total of 15 cells for each condition were analysed and the average plotted with the standard error of mean (SEM). Image intensity quantification was obtained by drawing cell outlines and determining the mean pixel intensity from the 16-bit images within ImageJ. A total of 37-126 cells were analysed per condition.

### E. coli cell death kinetics

*E. coli* JW2203 (OmpF-expressing) cells (25 ml) were grown in LB at 37 °C to OD_600_ ∼0.4 (∼10^8^ cells) prior to addition of chloramphenicol (20 µg/ml), to inhibit cell division, and either wild-type ColE9 or ColE9 K469C labelled with AF488, AF568 or AF647 (80 nM) added to the culture. A no-colicin control was also included. Colicin import was stopped at specific time points by removing 900 µl of culture and adding 100 μl of trypsin (1 mg/ml) and incubated at 37 °C for 30 min. 100 µl of each sample was diluted in LB (10^0^ -10^6^ fold), before plating (90 µl) on LB-agar plates (with appropriate antibiotics), and incubating for 16 hours at 37 °C. Colonies were counted and averaged from across triplicate plates. The number of CFUs were calculated using the following equation:

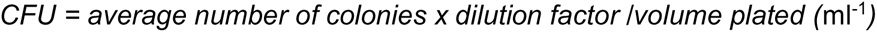

The natural log of each CFU value was plotted against time (mins), and the rate constant (k) of the reaction was calculated by representing the first order reaction as a straight line using equation: y = kx + c. The half-life of the reaction in cell killing kinetics are quoted as t_1/2_ values, where t_1/2_ = ln2/k.

