## Supplementary Figure for "Passive receptor dissociation driven by porin threading establishes active colicin transport through *Escherichia coli* OmpF"

### Supplementary Information

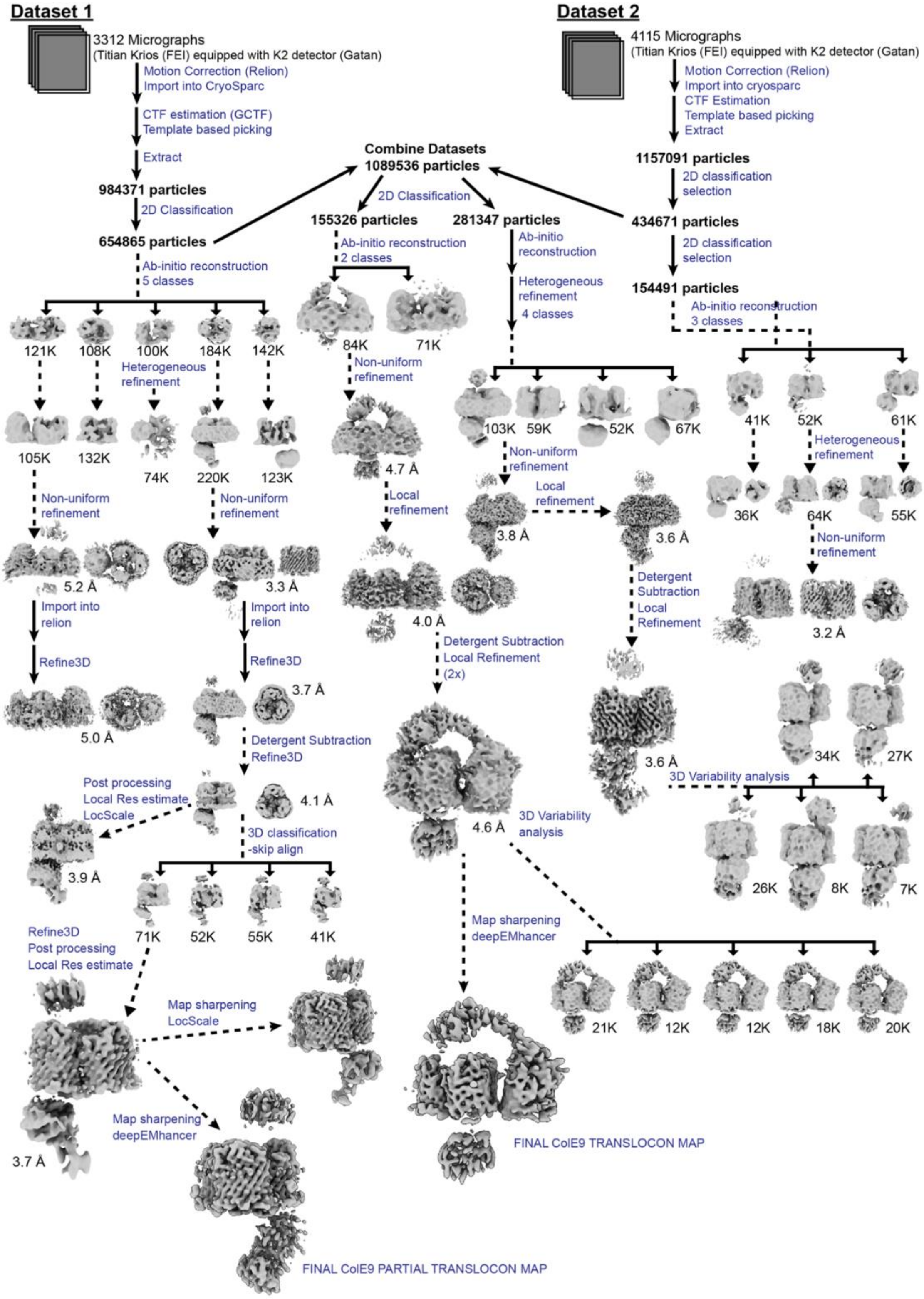


**Supplementary Figure 1**. Workflow for partial and full ColE9 translocon cryo-EM structures determination. DeepEMhancer sharpened final maps used for model building are indicated.

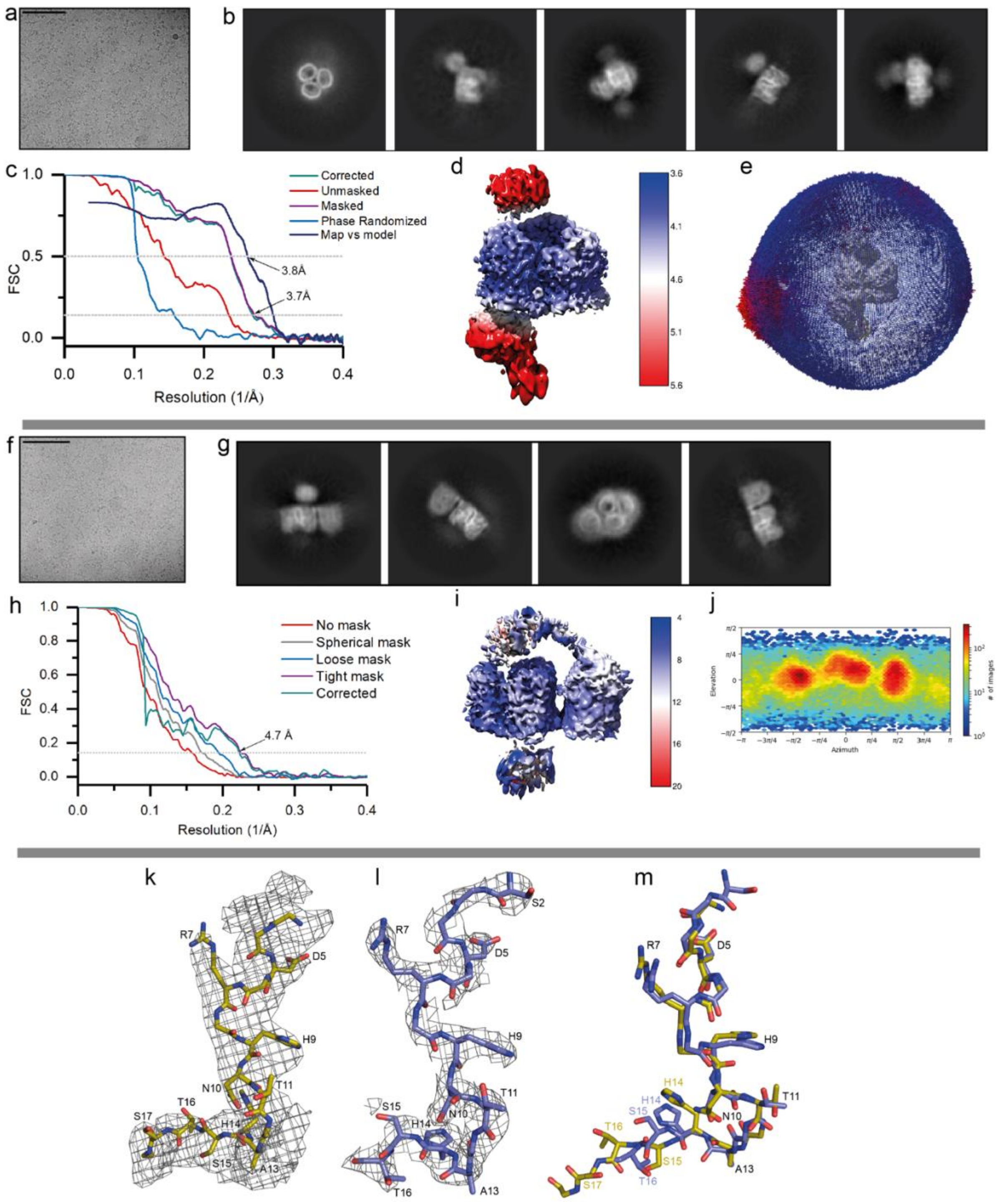


**Supplementary Figure 2. Single-particle cryo-EM analysis ColE9 translocon.** **a**, Representative cryo-EM image of ColE9 translocon (scale bar, 100 nm. **b**, Representative reference-free 2D class averages of particles selected for partial translocon map 3D reconstruction. **c**, Fourier Shell Correlation (FSC) plot generated after Relion postprocessing for the partial translocon map: curves for correlation between independently refined corrected (*teal*), unmasked (*red*), masked (*purple*), and phase-randomised (*blue*)

half-maps shown. In addition, this plot includes model-to-map FSC fit from Phenix refinement (*navy*). **d**, Partial translocon map coloured according to local resolution as estimated using Relion. **e**, Angular distribution plot from Relion refinement for partial translocon map (surface view in grey). **g**, Representative reference-free 2D class averages of particles selected for full translocon map 3D reconstruction. **h**, FSC plot showing curves for correlation between independently refined half-maps with no mask (*red*), spherical mask (*grey*), loose mask (*blue*), tight mask (*purple*) and corrected (*teal*). **i**, Full translocon map coloured by local resolution estimate as implemented in cryoSPARC. **j**, Angular distribution of particles contributing to the final full translocon map. **k**, Cryo-EM partial translocon map density (*grey mesh*) is shown at threshold level of 3.0 surrounding OBS1 (*gold sticks*). Figure generated in pymol. **l**, X-ray crystallography electron density map (*grey mesh*) is shown at a threshold level of 1.0 surrounding OBS1 (*purple*) as observed in the OmpF-OBS1 peptide complex (PDB ID, 3O0E). **m**, Overlay of OBS1 (*gold*) from the partial translocon model and the OBS1 peptide (*purple*) from the crystal structure of the OmpF-OBS1 peptide complex, showing the near-identical conformation of the peptide adopted in both structures. Residues are denoted and labelled according to stick colour.

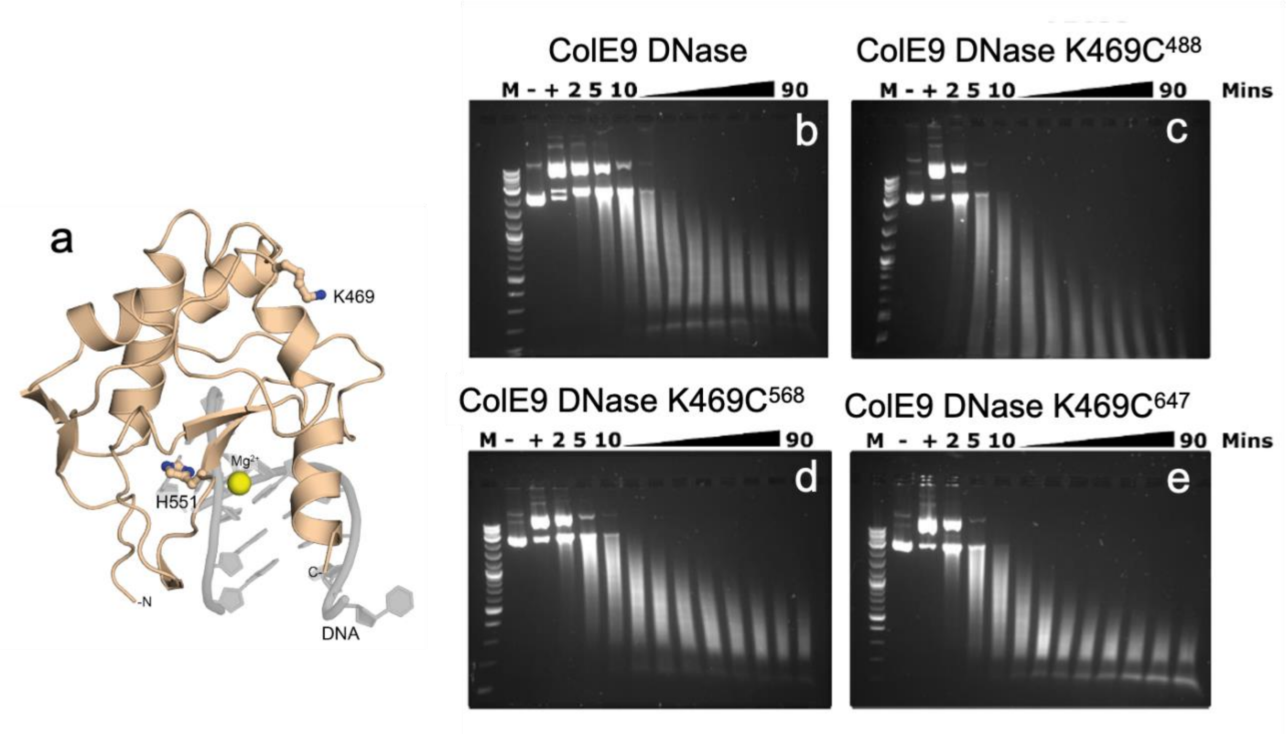


**Supplementary figure 3. Attachment of AlexaFluor (AF) dyes to ColE9 DNase K469C has no effect on endonuclease activity**. **a**, Crystal structure of the ColE9 DNase domain (*cream*) bound to dsDNA (*grey*) and catalytic divalent cation (PDB ID 1V14) showing the position of Lys469 on the periphery of the DNA binding site. Also shown is His551, a catalytic residue that was mutated to alanine in order to inactivate the toxin in fluorescence transport assays (denoted throughout as ColE9*). **b – e**, Plasmid nicking assays of 1 µg/ml pUC19 DNA in 50 mM TEA buffer, pH 7.5 containing 20 mM $MgCl_2$, at room temperature to which 200 nM ColE9 DNase domain was added, labelled at K469C with different fluorophores (AF488, AF568 and AF647). Samples of each reaction were taken over 90 minutes and separated by agarose gel and DNA stained with SYBR Safe (Invitrogen). *M*, marker lane. -, substrate DNA. +, substrate DNA plus designated DNase domain at the start of the reaction. The rate of DNA hydrolysis was the same for wild-type ColE9 DNase domain and the DNase carrying different fluorophores at K469C.

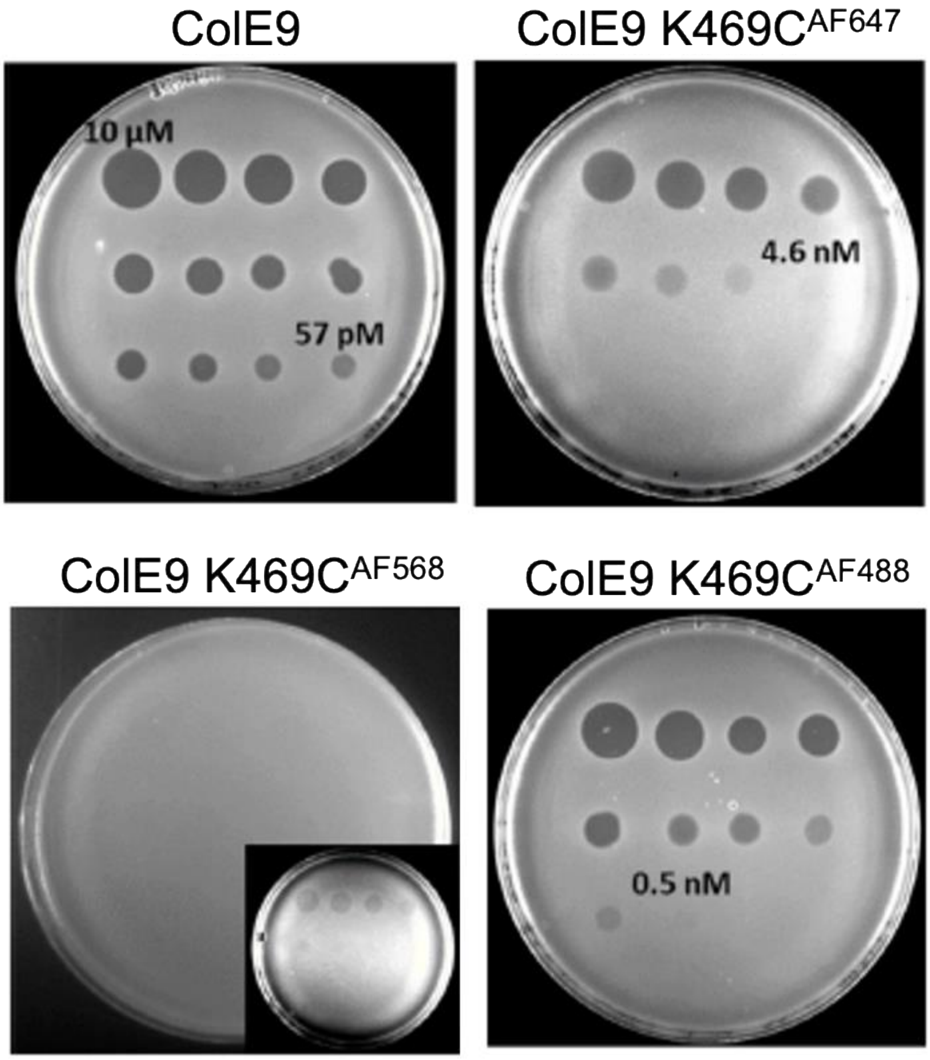


**Supplementary figure 4. Covalent attachment of AF dyes to ColE9 K469C modulates colicin-mediated killing of *E. coli* JM83 cells.** Panel shows plate killing assays comparing wild-type ColE9 cytotoxicity against ColE9 K469C labelled with AF488, AF568 and AF647 against *E. coli* JM83 cells, which express both *ompF* and *ompC*. A serial dilution of colicin (10 µM to 57 pM in 3-fold serial dilutions) was spotted onto each plate and incubated overnight at 37 °C. Zones of clearance show colicin induced cell-killing. ColE9 K469C$^{AF488}$ and ColE9 K469C$^{AF647}$ had reduced killing relative to wild-type ColE9. By contrast, ColE9 K469C$^{AF568}$ little toxic activity; an enhanced imaged is shown in the inset to indicate the presence of feint killing zones.

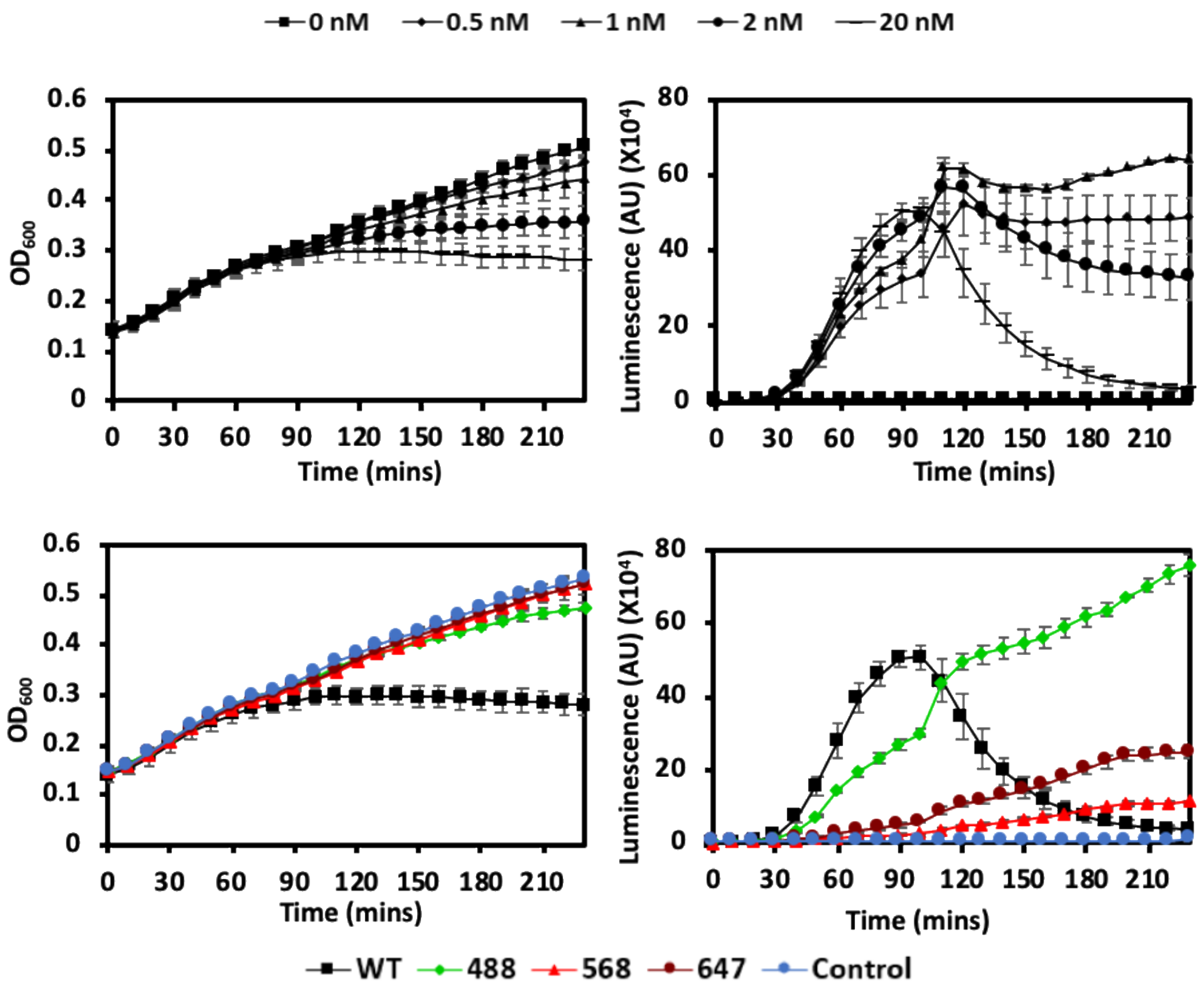


**Supplementary figure 5. AF labels attached to ColE9 K469C slow activation of the SOS response in *E. coli*.** Panels a and c show growth curves and panels b and d show *lux* bioluminescence emission data over time, for the strain *E. coli* DPD1718 grown in LB media at 37 °C and treated with wild-type ColE9 or ColE9 K469C labelled with AF488, AF568 or AF647. *E. coli* DPD1718 contains a fusion of the LexA regulated *recA* promoter region of the SOS operon of *E. coli* upstream of the *P. luminescens luxCDABE* operon. ColE9 DNase entering the cytoplasm nicks the chromosome and activates the SOS response, which is quantitated through *lux* bioluminescence. **a**, *E. coli* DPD1718 cells were incubated with increasing concentrations of wild-type ColE9 (0.5 - 20 nM) and growth of cells monitored at $OD_{600}$ in a plate reader. Growth was completely inhibited by 20 nM ColE9. **b**, *E. coli* DPD1718 bioluminescence emission data appears after 30 min following treatment with the same concentration range of ColE9 as in panel a. Low concentrations of ColE9

(0.5 - 2 nM) activate the SOS response but the signal plateaus as cell growth begins to outcompete cell killing, as reported previously. At 20 nM, all cells are killed hence the drop in bioluminescence signal after the peak at ~90 min, which coincides with the cessation of growth. Error bars reflect SEM from 3 experiments. **c**, Growth curves for *E. coli* DPD1718 untreated (*blue circles*) and treated with 20 nM ColE9 (*black squares*) or ColE9 K469C carrying different AF dyes; AF488 (*green diamond*s), AF568 (*red triangles*), AF647 (*maroon circles*). With the exception of AF488, which shows mild toxicity, the other fluorophores suppress cell killing by ColE9 in this assay. **d**, *E. coli* DPD1718 bioluminescence emission data following addition of ColE9 K469C (20 nM) labelled with different AF dyes. Regardless of their poor killing profiles in this strain, all AF-labelled ColE9s activated the SOS response, from which we conclude all reach the cytoplasm to cleave DNA. AF568 had the greatest inhibitory effect on SOS activation by ColE9 in this assay followed by AF647 and then AF488. The continual and steady increase in lux bioluminescence is consistent with cell growth outcompeting cell killing by these colicin constructs. Error bars reflect SEM from 3 experiments.

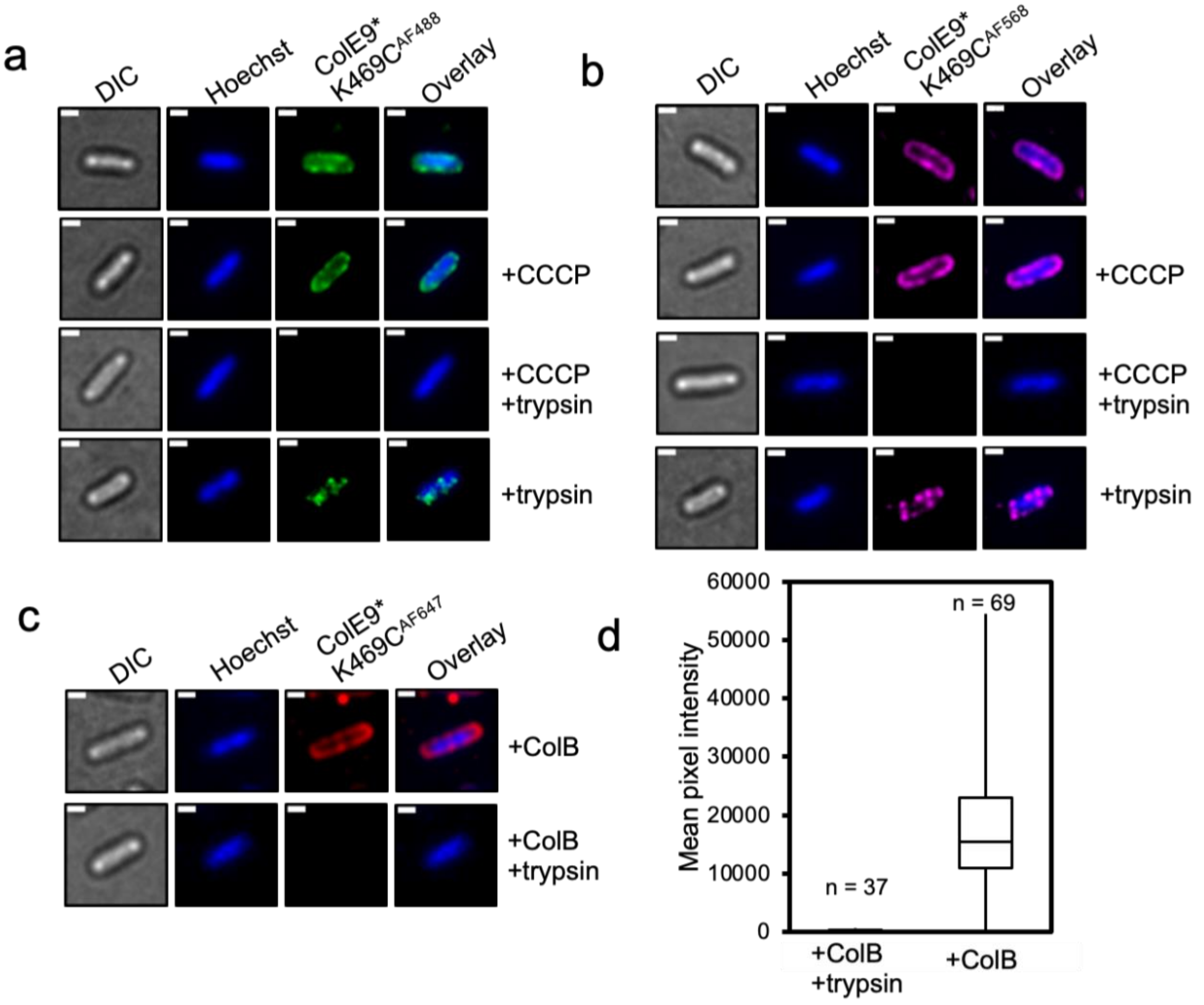


**Supplementary figure 6. Import of ColE9 K469C labelled with different fluorophores is PMF-dependent**. **a,** Widefield fluorescence microscopy images of *E. coli* JM83 cells labelled with ColE9* K469C$^{AF488}$ (1.5 µM) and Hoechst stain (20 µM), for 30 mins at 37 °C with or without trypsin treatment with or without prior treatment with CCCP. **b,** Repetition of experiment in panel a, with ColE9* K469C$^{AF568}$. For panels a and b, every set of horizontal panels shows the same cell in DIC (*grey*), Hoechst DNA stain (*blue*) and with ColE9* K469C$^{AF488}$ fluorescence (*green*) or ColE9* K469C$^{AF568}$ fluorescence (*pink*) and overlays of DNA and fluorescent colicins are also shown. In panels a and b, destruction of the PMF followed by trypsin treatment removed all surface-bound fluorescent signal. Single treatment with either CCCP or trypsin alone, showed retention of fluorescence signal at the cell periphery and internally, respectively. Taken together, AF labelled ColE9 is only internalised with a functional PMF. **c**, Widefield fluorescence microscopy images of *E. coli*

JM83 cells labelled with ColE9* K469C$^{AF647}$ (1.5 µM) and Hoechst stain (20 µM) with or without prior treatment with colicin B (ColB, 2.5 µM). Subsequent treatment with trypsin was used to remove surface-bound molecules. ColB is a pore-forming toxin that depolarizes the cytoplasmic membrane. Each panel shows the same cell seen in DIC (*grey*), Hoechst DNA stain (*blue*) and ColE9* K469C$^{AF647}$ fluorescence (*red*). ColE9* K469C$^{AF647}$ remained bound to the OM in the presence of ColB, but fluorescence was lost after treatment of these cells with trypsin. Scale bars, 1 µm. **d**, Box and whisker plots of fluorescence microscopy data for *E. coli* JM83 cells treated with ColB and labelled with ColE9* K469C$^{AF647}$ with and without trypsin treatment: whiskers represent minimum and maximum mean pixel intensity, box shows 1$^{st}$ and 3$^{rd}$ quartile with the median shown as a line. Microscopy data were collected as in c. The mean pixel intensity of ColE9* K469C$^{AF647}$ per cell was measured for each cell condition. *n*, number of cells used from two biological replicates in each case. The data show that prior treatment of cells with ColB prevents the translocation of ColE9* K469C$^{AF647}$ across the OM, similar to the effects of CCCP (see **Figure 5**).

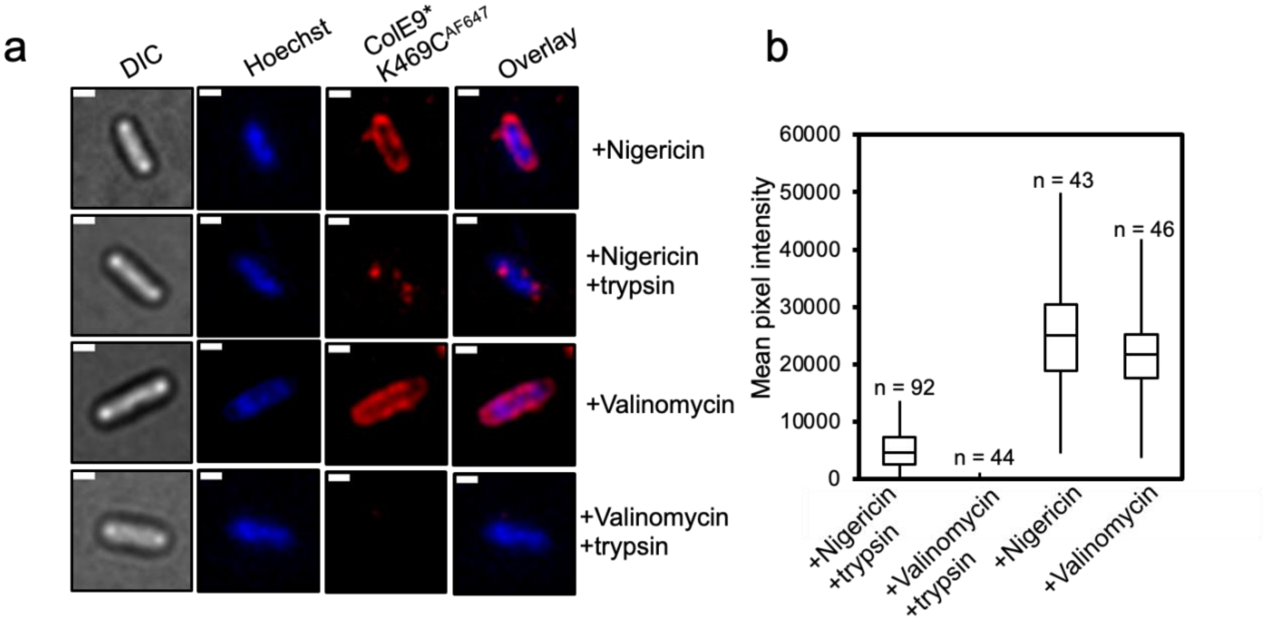


**Supplementary figure 7**. **The electrical potential of the PMF drives ColE9* K469C$^{AF647}$ import in *E. coli*.** **a,** Widefield fluorescence microscopy images of *E. coli* JM83 cells treated with nigericin or valinomycin (5 µM) and then labelled with ColE9* K469C$^{AF647}$ (1.5 µM) and Hoechst stain (20 µM) for 30 min at 37 $^{o}$C with or without trypsin treatment. Each panel shows the same cell in DIC (*grey*), Hoechst DNA stain (*blue*) and ColE9* K469C$^{AF647}$ fluorescence (*red*). Overlays of DNA and ColE9* K469C$^{AF647}$ fluorescence are also shown. ColE9* K469C$^{AF647}$ remained bound to the OM in the presence of both nigericin and valinomycin, but fluorescence was lost after trypsin treatment in valinomycin-treated cells. Scale bar, 1 µm. **b**, Box and whisker plots of fluorescence microscopy data for *E. coli* JM83 cells treated with nigericin or valinomycin and labelled with ColE9* K469C$^{AF647}$ with and without trypsin treatment: whiskers represent minimum and maximum mean pixel intensity, box shows 1$^{st}$ and 3$^{rd}$ quartile with the median shown as a line. Microscopy data were collected as in panel a. The mean pixel intensity of ColE9* K469C$^{AF647}$ per cell was measured for each cell condition. *n*, number of cells used across two or three biological replicates. The data show that when valinomycin destroys the electrical potential ($\Delta\Psi$) of the PMF that translocation of ColE9* K469C$^{AF647}$ across the OM is inhibited.

**Supplementary Table 1. Cryo-EM data collection and processing statistics**

| | Full ColE9 Translocon<br>EMDB-12577<br>PDB-7NSU | Partial ColE9 Translocon<br>EMDB-12576<br>PDB-7NST |
|---|---|---|
| **Data collection** | | |
| Microscope | Titan Krios 3Gi | Titan Krios 3Gi |
| Voltage (kV) | 300 | 300 |
| Detector | K2 | K2 |
| Recording mode | counting | counting |
| Magnification | 130,000 | 130,000 |
| Movie/micrograph pixel size (Å) | 1.047 | 1.047 |
| Dose rate (e-/Å$^2$/sec) | 4.9 | 4.997 |
| Number of frames per movie | 50 | 50 |
| Movie exposure time (s) | 10 | 10 |
| Total dose (e-/Å$^2$) | 49 | 50 |
| Defocus range (um) | -1.5 to -3.5 | -1.5 to -3.5 |
| **EM data processing** | | |
| Number of movies/micrographs | 7427 | 3312 |
| Box size (px) | 280 | 280 |
| Particle number (total) | 1,089,536 | 984,371 |
| Particle number (post 2D) | 155,326 | 654,865 |
| Particle number (used in final map) | 83,697 | 71,190 |
| Symmetry | C1 | C1 |
| Map resolution (FSC 0.143) | 4.6 | 3.7 |
| Local resolution range (FSC 0.5) | 4.6-16 | 3.6-9.2 |
| Map sharpening B-factor (Å$^2$) | -203 | -185 |
| **Model Building and Validation** | | |
| Initial model used | PBD-7NST(partial), TolB-TBE 2IVZ, BtuB-ColE3 2YSU, ColE9 5EW5 | TolB-TBE 2IVZ, ColE9 5EW5, ompF 3K19 |
| Model composition | | |
| Non-hydrogen protein atoms | 36210 | 25304 |
| Protein residues | 2414 | 1706 |
| Nucleotides (RNA) | 0 | 0 |
| RMSD from ideal | | |
| Bond length (Å) | 0.003 | 0.003 |
| Bond angles (°) | 0.699 | 0.615 |
| Validation | | |
| Molprobity score | 2.34 | 1.66 |
| Clashscore | 10.9 | 8.42 |
| Rotamers outliers (%) | 3.75 | 0.89 |
| FSC (0.5) model-vs-map | 8.3 | 3.9 |
| CC model-vs-map (masked) | 0.49 | 0.8 |

| | | |
|---|---|---|
| Ramachandran plot | | |
| Favored (%) | 95.03 | 96.75 |
| Allowed (%) | 4.85 | 3.2 |
| Outliers (%) | 0.13 | 0.06 |

**Supplementary Table 2. *E.coli* strains used in this study**

| Strain | Genotype | References |
|---|---|---|
| BL21 (DE3) | $F^{-}$ *ompT hsdS*$_{B}$($r_{B}^{-}$ $m_{B}^{-}$) *gal dcm* (DE3) | Novagen |
| JM83 | rpsL ara Δ(lac-proAB) φ80dlacZΔM15 | (Yanisch-Perron et al., 1985) |
| RK5016 | Δ*lac*(*U169*) *araD139 rpsL gyrA thi non metE argH recA* Δ*btuB* | (Heller et al., 1985) |
| DPD1718 | *lac* Cam$^{r}$ *lacZ*::*recA* φ*luxCDABE* | (Davidov et al., 2000) |
| JW2203 | F-, *Δ(araD-araB)567*, *ΔlacZ4787*(::rrnB-3), *λ*$^{-}$, *ΔompC768::kan*, *rph-1*, *Δ(rhaD-rhaB)568*, *hsdR514* | Keio<br>(Baba et al., 2006) |
| JW0912 | F-, *Δ(araD-araB)567*, *ΔlacZ4787*(::rrnB-3), *λ*$^{-}$, *ΔompF746::kan*, *rph-1*, *Δ(rhaD-rhaB)568*, *hsdR514* | Keio<br>(Baba et al., 2006) |
| 113-3<br>(DSM-1900) | *Δ* metE mutant of wild-type *E.coli* (ATCC 9637) | (Davis and Mingioli, 1950) |

**Supplementary Table 3. Description of plasmids used in this study**

| Plasmid name | Plasmid description | References |
|---|---|---|
| pUC19 | | NEB |
| pCS4 | pET21a-ColE9-Im9 | (Garinot-Schneider et al., 1997) |
| pMLF07 | pET21a-ColE9-K469C-Im9 | This work |
| pMLF08 | pET21a-ColE9-H551A-K469C-Im9 | This work |
| pRJ353 | pET21d-ColE9-DNase-Im9 | (Garinot-Schneider et al., 1996a) |
| pBC1 | pET21d-ColE9-DNase-K469C-Im9 | This work |
| pNGH136 | pET21a-A33C ColE9 T-R-$TEV_{His6}$ | (Housden et al., 2013) |
| pNGH89 | pET21d-P201C $TolB_{His6}$ | (Housden et al., 2013) |
| pEE01 | pACYCDuet-1$ColB_{His6}$ | This work |
| pNGH96 | pET21a-ColE9-W39A-Im9 | Housden et al. 2010 |
| pREN151 | pET21a-ColE9-343-418 | This work |
